# The coevolution of sex-specific dominance and allelic diversity at sexually antagonistic loci

**DOI:** 10.1101/2024.05.06.592709

**Authors:** Mattias Siljestam, Claus Rueffler, Göran Arnqvist

## Abstract

Sexually antagonistic (SA) selection can promote genetic diversity. Current theory predicts biallelic polymorphism in SA loci primarily under strong selection or dominance reversal between the sexes. Yet, selection is often weak, several candidate SA loci harbour more than two alleles, and the prevalence of dominance reversal remains unclear. We present a model in which allelic effects at a quantitative-trait locus coevolve with sex-specific promoter affinities that determine allelic dominance, under distinct female and male phenotypic optima. We show that gradual coevolution between allelic values and promoter affinities can generate a positive feedback between beneficial sex-specific dominance and allelic divergence. Importantly, this feedback does not require an established balanced polymorphism, but can be initiated during selective sweeps, during evolutionarily transient protected polymorphism, or from variation maintained by mutation– selection–drift balance. Under weak or asymmetric selection, repeated diversification can generate polyallelic polymorphism, with several alleles forming dominance hierarchies that reduce segregation load. These results hold for both autosomal and X-linked loci and do not require complete dominance reversal.

## Introduction

**O**ur understanding of the maintenance of genetic variation in loci and traits under selection is still incomplete, and the relative roles of mutation-selection balance, balancing selection and other processes remain debated. Quantitative assessments suggest that some form of balancing selection must contribute to the maintenance of polymorphisms for fitness-related traits (Charlesworth and Hughes, 2000; Mitchell-Olds et al., 2007; Charlesworth, 2015). A potential generator of balancing selection are trade-offs resulting in antagonistic pleiotropy, whereby alternative alleles are favoured by selection in different contexts (e.g., environmental conditions, ontogenetic stages, age classes or sexes). Previous theory suggests that the conditions under which antagonistic pleiotropy maintains biallelic polymorphism (BAP) in SA loci may be quite restricted (Prout, 2000; Flintham et al., 2026). In particular, under additive allelic effects, the conditions for stable polymorphism become increasingly restrictive under weak or asymmetric selection (e.g., Kidwell et al., 1977; Flintham et al., 2025). Polymorphism can instead be promoted by, for example, disassortative mating by genotype (Arnqvist, 2011), dominance under sex-linkage (Rice, 1984; Patten and Haig, 2009) or dominance reversal (Kidwell et al., 1977; Fry, 2010). Dominance reversal, where each allele shows dominance in the context in which it is beneficial and recessivity when it is disfavoured, has a particularly high potential to promote the maintenance of genetic variation (Kidwell et al., 1977; Wilder et al., 2016; Wittmann et al., 2017; Grieshop and Arnqvist, 2018; Connallon and Chenoweth, 2019; Grieshop et al., 2024; Khudiakova et al., 2025).

Dominance reversal has traditionally been considered unlikely (Hedrick, 1999; Prout, 2000), partly based on the observation that mutations with major effects (e.g., causing disorders in humans) tend to be unconditionally dominant or recessive (Curtsinger et al., 1994). Although our understanding of the proximate mechanisms of dominance is still limited (Huber et al., 2018), both theory (Otto and Bourguet, 1999; Spencer and Priest, 2016) and data (Billiard and Castric, 2011) suggest that dominance relationships can and do evolve (Grieshop et al., 2024). Specifically, existing theory (reviewed in Otto and Bourguet, 1999) shows that dominance may readily evolve when allelic polymorphism is maintained by balancing selection, but that this is less likely during selective sweeps or in polymorphisms maintained under mutation-selection-drift balance. Although empirical data on dominance reversal is still limited, a few studies have provided evidence for dominance reversal across the sexes (Meiklejohn et al., 2014; Barson et al., 2015; Grieshop and Arnqvist, 2018; Pearse et al., 2019; Mérot et al., 2020; Geeta Arun et al., 2021; Puixeu et al., 2023; Kaufmann et al., 2024) and environments (Posavi et al., 2014; Chen et al., 2015; Karageorgi et al., 2024).

Sexually antagonistic (SA) selection is a potentially near-ubiquitous generator of antagonistic pleiotropy in taxa with separate sexes, because the same allelic effects are repeatedly exposed to two distinct and persistent genetic environments (males and females) in which selection can act in opposing directions (Bonduriansky and Chenoweth, 2009). Both sexes contribute to the gene pool of the next generation through their gametes, so the resulting evolutionary outcome reflects the balance between these opposing sex-specific selection gradients. Explicit genetic models have shown that sex-specific dominance is a predicted outcome in SA loci (Fry, 2010; Spencer and Priest, 2016; Connallon and Chenoweth, 2019) and a few empirical studies have confirmed that SA alleles beneficial in one sex indeed tend to be dominant in that sex but recessive in the other (Barson et al., 2015; Grieshop and Arnqvist, 2018; Pearse et al., 2019; Glaser-Schmitt et al., 2021).

A notable and intriguing property of several putative SA loci is the apparent segregation of more than two alleles. Polyallelic polymorphism (PAP) has, for example, been documented for the VGLL3 locus in salmon (Barson et al., 2015; Sinclair-Waters et al., 2022), the Cyp6g1 locus in *Drosophila melanogaster* (Schmidt et al., 2010; Hawkes et al., 2016), an X-linked regulatory element in *D. melanogaster* (Glaser-Schmitt and Parsch, 2018; Glaser-Schmitt et al., 2021), the DsFAR2 locus in *D. serrata* (Rusuwa et al., 2022), the Pgi locus in butterflies (Niitepõld and Saastamoinen, 2017) and neurogenetic loci in voles (Lonn et al., 2017). The maintenance of PAP in loci under selection is generally not well understood (Lewontin et al., 1978; Spencer and Mitchell, 2016), and the presence of PAP in candidate SA loci suggests that additional theory is required. We note that sexual conflict mediated by the compatibility between a pair of interacting loci with sex-limited expression (i.e., interlocus sexual conflict; Arnqvist and Rowe, 2005) can act to maintain PAP through specific pairwise compatibility between, for example, male ligands and female receptors (Gavrilets and Waxman, 2002; Haygood, 2004), but this is not directly applicable to SA loci.

In this study, we develop a mathematical model to assess under which conditions one should expect the emergence of sex-specific dominance through gradual evolution acting at a locus affecting a quantitative trait under SA selection and how this, in turn, affects the evolution of allelic polymorphism at that locus. Additionally, given that most previous models of SA loci are restricted to biallelic scenarios, we investigate whether the evolution of sex-specific dominance can lead to the origin and maintenance of PAP in SA loci. We show that gradual coevolution between allelic values and sex-specific promoter affinities can generate a positive feedback between beneficial sex-specific dominance and allelic divergence. Importantly, this feedback is not restricted to cases where polymorphism is already maintained by balancing selection: it can be initiated during selective sweeps, during evolutionarily transient protected polymorphism, or from genetic variation maintained by mutation–selection– drift balance. Under weak or asymmetric selection, where evolutionarily stable polymorphism is not expected in the absence of beneficial dominance, this coevolution can ultimately generate PAP and thereby reduce segregation load further. We arrive at qualitatively similar results for dominance effects in X-linked loci.

### The model

We consider allelic evolution at a locus subject to sexually antagonistic (SA) selection in a panmictic population of constant size with non-overlapping generations (Fisher, 1930; Wright, 1931). The locus affects a quantitative trait, denoted *z*, for which females and males have different phenotypic optima. The trait *z* could represent any shared characteristic, such as morphology, colouration, or a life-history trait.

The fitness (viability) *w* of an individual decreases according to a Gaussian function with increasing deviation of its phenotypic trait value *z* from the sex-specific optimum, as illustrated in Figure 1. Without loss of generality, we assume that the female optimum is smaller than the male optimum (Fig. 1). The widths of the Gaussian functions (*σ*_♀_ and *σ*_♂_) determine the strength of selection in each sex, while their widths relative to the distance *δ* between the optima determine the strength of sexually antagonistic selection. Specifically, larger values of 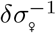 and 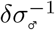 correspond to stronger selection relative to the distance between the sex-specific optima. For presentation purposes, we set *δ* = 1 (with sex-specific optima *−*0.5 for females and 0.5 for males) and therefore use 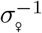 and 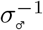 as measures of selection strength throughout the main text.

**Figure 1.**
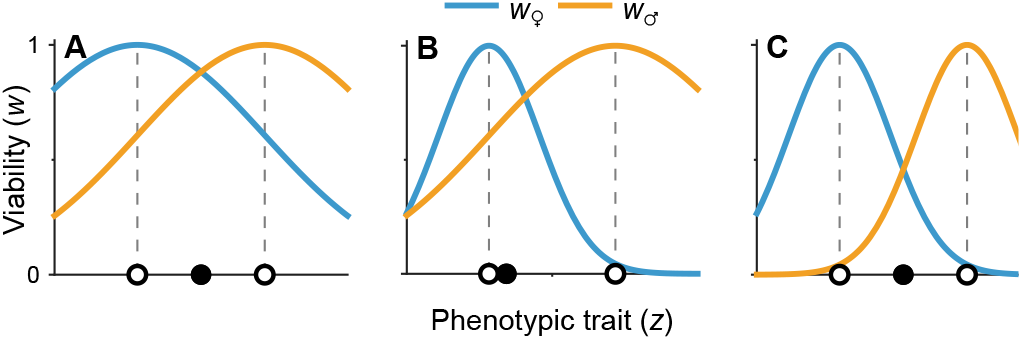
Sex-specific fitness (viability) *w* as a function of the phenotypic trait *z*. Open dots indicate the phenotypic optima for females and males, while filled dots indicate the trait value that maximizes geometric mean fitness across the two sexes. In **A**, both sexes are under weak selection 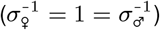, while in **B** females are under strong selection while males are under weak selection 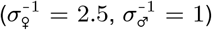. In **C**, both sexes are under strong selection 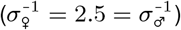.

We start by considering an autosomal locus and later extend the model to an X-linked locus. Each allele is characterized by an allelic value *x*_*i*_, which influences the phenotypic trait *z*. Thus, *z* is a function of the allelic values of the two alleles *x*_*i*_ and *x*_*j*_ carried at the locus, *z*(*x*_*i*_, *x*_*j*_). We assume no parental effects, so that the order of the arguments is irrelevant, *z*(*x*_*i*_, *x*_*j*_) = *z*(*x*_*j*_, *x*_*i*_). For individuals homozygous for allele *x*_*i*_, the phenotype equals the allelic value, *z*(*x*_*i*_, *x*_*i*_) = *x*_*i*_. For heterozygotes carrying alleles *x*_*i*_ and *x*_*j*_, the resulting phenotype *z* falls between *x*_*i*_ and *x*_*j*_, with the precise value determined by dominance.

To allow for the evolution of sex-specific dominance, we employ the promoter affinity model of Van Dooren (1999) (see Flintham (2025) for a recent sex-specific implementation). Each allele is, in addition to its allelic value *x*_*i*_, defined by an upstream promoter sequence, having affinities *α*_*i*, ♀_ and *α*_*i*, ♂_, interpreted as the allele’s relative recruitment of transcriptional machinery in the two sexes. Under a fixed total transcriptional output, these affinities determine the proportion of gene product contributed by each allele, yielding the heterozygote phenotype

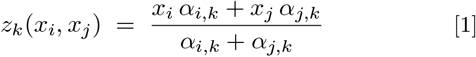

(see Eq. 1 in Van Dooren, 1999). Defining

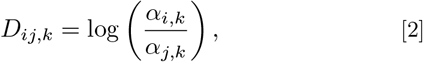

as the difference in log-affinity between the two alleles in sex *k*, Equation 1 can be written as

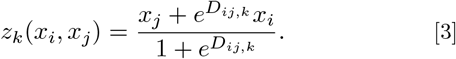

The corresponding (sex-specific) dominance weight on allele *i* is 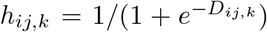, such that *z*_*k*_ = *h*_*ij,k*_*x*_*i*_ + (1 *−h*_*ij,k*_)*x*_*j*_.

Equal promoter affinities, *α*_*i,k*_ = *α*_*j,k*_ (or *D*_*ij,k*_ = 0), correspond to additivity (Fig. 2**A**), while *α*_*i,k*_*≫ α*_*j,k*_ makes allele *i* nearly completely dominant in sex *k* (*h*_*ij,k*_ *≈* 1). Dominance is sex-specific whenever *h*_*ij*, ♀_ ≠ *h*_*ij*, ♂_ (Fig. 2**B**–**C**), which occurs whenever the affinity ratios differ between the sexes (i.e., *α*_*j*, ♀_*/α*_*i*, ♀ ≠_*α*_*j*, ♂_ */α*_*i*, ♂_), and can yield dominance reversals across the sexes when an allele is (partly) dominant in females (*α*_*i*, ♀_ *> α*_*j*, ♀_) but (partly) recessive in males (*α*_*i*, ♂_ *< α*_*j*, ♂_; Fig. 2**C**), or vice versa. Note that *α*-values have no effect in homozygous individuals. Therefore, promoter affinities can only be under selection in populations with more than one segregating *x*-allele.

**Figure 2.**
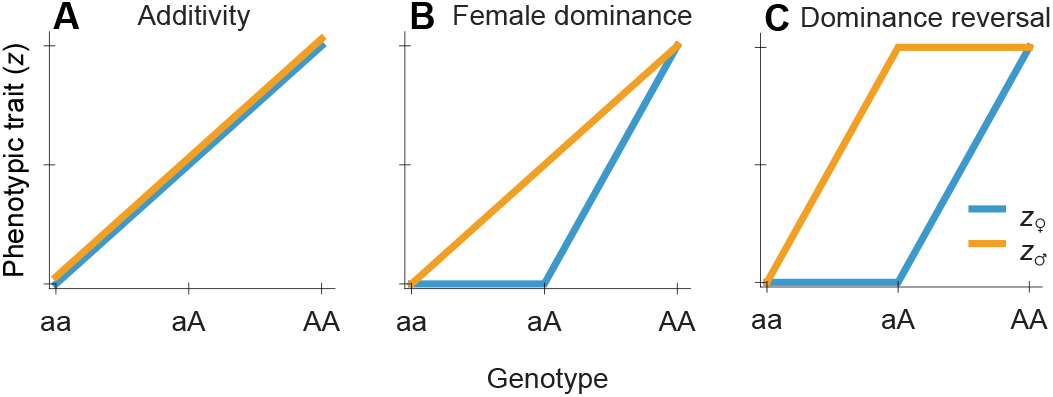
Illustrations of phenotypic trait values resulting from two alleles, one favoured in females (*x*_*a*_) and one in males (*x*_*A*_). **A.** Additive allelic effects obtain when the two alleles have equal promoter affinities within both sexes (*α*_*a*,♀_ = *α*_*A*,♀_ and *α*_*a*,♂_ = *α*_*A*,♂_), corresponding to *h* = 0.5. **B**. Sex-specific dominance with dominance in females (*α*_*a*,♀_ *≫ α* _*A*,♀_ *→ h* = 0) and additivity in males (*α*_*a*,♂_ = *α*_*A*,♂_ *→ h* = 0.5). **C**. Dominance reversal (*α*_*a*,♀_ *≫ α*_*A*, ♀_ *→ h* = 0 and *α*_*a*, ♂_ *≪ α*_*A*,♂_ *→ h* = 1).

While classic theory predicts that the X-chromosome should be enriched with SA loci (Rice, 1984), more recent work has shown that this is not necessarily the case (Fry, 2010). Some empirical data are consistent with an important role for X-linked sites in sex-specific traits (Reinhold, 1998; Gibson et al., 2002; Dean and Mank, 2014) but testing for an enrichment of SA loci in X is challenging (Ruzicka and Connallon, 2020). Here, we analyse a version of our model for X-linked loci. This version follows the same principles, with the exception that dominance can only be expressed by females, while a male that is hemizygous for *x*_*i*_ has a phenotype directly determined by its allelic value, *z* = *x*_*i*_, as would for example be the case under complete dosage compensation. Consequently, dominance in an X-linked locus inherently exhibits sex-specific characteristics, and we demonstrate that the outcomes are qualitatively similar to those observed with sex-specific dominance in an autosomal locus.

### Analysis

#### Invasion analysis

We use evolutionary invasion analysis (Metz et al., 1992; Geritz et al., 1998; Kisdi and Geritz, 1999; Doebeli, 2011) to study evolution of allelic values *x* and sex-specific promoter affinities *α*. This approach is based on invasion fitness, the long-term growth rate of rare mutant alleles of small effect. For details, see Section S3 in the Supplementary material. Based on this approach, we derive conditions for evolutionary branching at an allelic value *x*\* where directional selection ceases to act. If the branching condition is fulfilled, a monomorphic allelic lineage splits into two lineages that subsequently diverge due to disruptive selection.

#### Simulations

We further investigate allelic diversification using individual-based computer simulations based on Wright–Fisher population dynamics with mutation and selection (Fisher, 1930; Wright, 1931). At the trait locus *x*, mutational effects are drawn from a normal distribution with an expected step size of 0.08. The sex-specific optima (Fig. 1) are separated by one unit (*δ* = 1), equivalent to ~13 average mutational steps. Unless stated otherwise, simulations are initialized with a monomorphic population one unit from *x*\*, ensuring comparable initial directional selection across simulations.

We use a conservative per-gene mutation rate of *µ*_*x*_ = 5 *×*10^*−*6^ for mutations affecting the modelled trait. For comparison, assuming a per-base mutation rate of *µ≈* 1*×*10^*−*8^ (Bergeron et al., 2023), this corresponds to an effective mutational target of only ~ 500 bp per gene, excluding substitutions that do not affect the trait. We use a population size of *N* = 5 *×*10^4^, giving a mutational input of 2*Nµ*_*x*_ = 0.5, corresponding to an average of one new mutant allele affecting *x* every two generations. This mutational input is conservative relative to many natural populations, particularly in taxa with larger effective population sizes (Lynch et al., 2023). Moreover, mutational processes beyond exonic base substitutions may further increase the effective per-gene mutation rate. In Figure 3**iic** and the right column of Figure 5, we instead use *N* = 5 *×*10^3^, reducing the mutational input to 2*Nµ*_*x*_ = 0.05.

**Figure 3.**
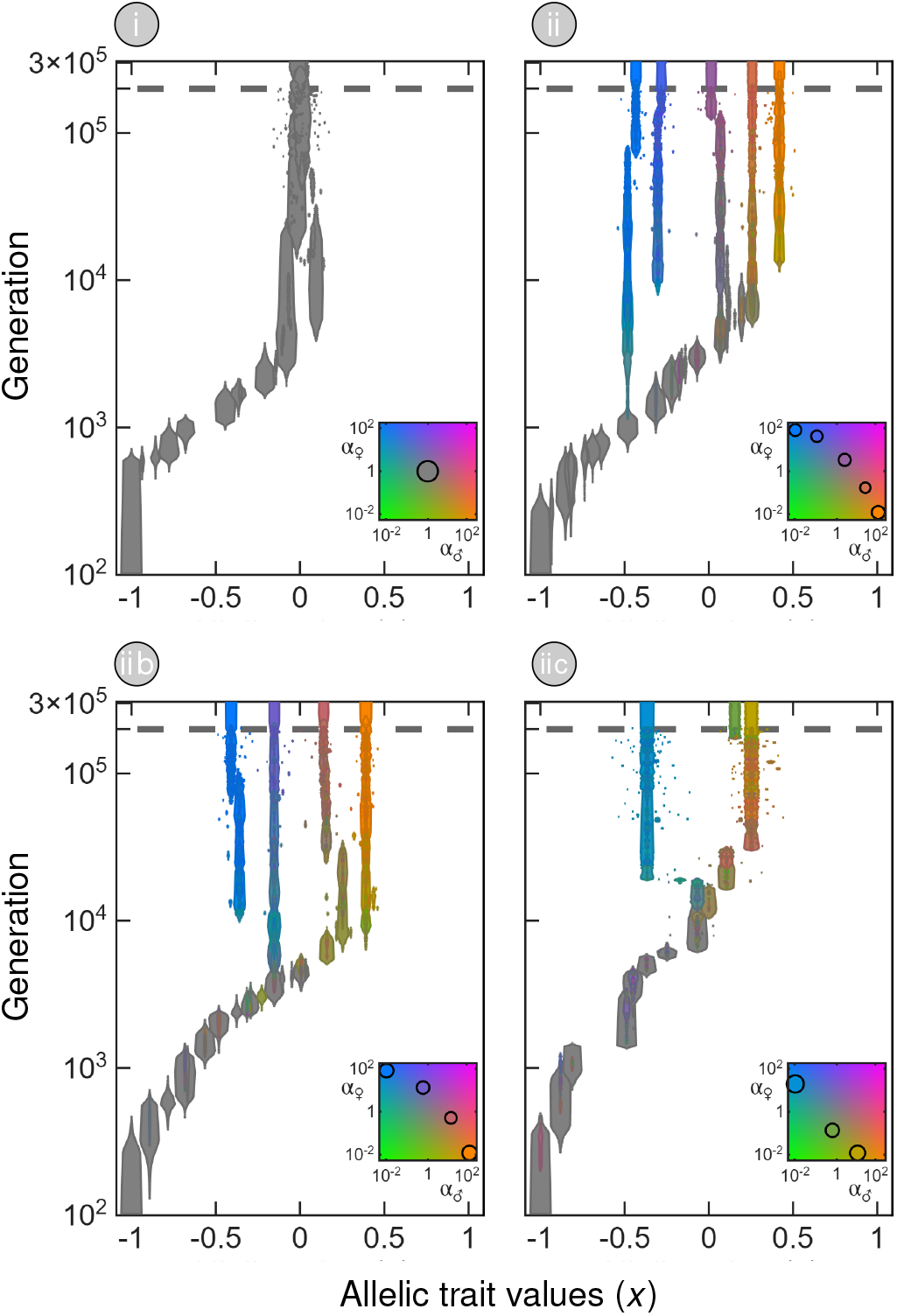
Individual-based simulations illustrating three routes by which the coevolutionary feedback between beneficial sex-specific dominance and allelic divergence can be initiated under weak selection 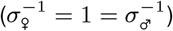. In all simulations, allelic evolution approaches the evolutionary attractor *x*\* = 0. In panel **i**, promoter affinities are fixed, and selective sweeps are followed by an evolutionarily transient protected polymorphism between alleles on opposite sides of *x*\* that persists for approximately 10^5^ generations. With evolving promoter affinities, the coevolutionary feedback is initiated during a selective sweep in panel **ii**, during a transient protected polymorphism around *x*\* in **iib**, and from variation maintained by mutation–selection–drift balance after reaching *x*\* in **iic.** Panel **iic** uses a smaller population size (*N* = 5 *×*10^3^; *N* = 5 *×*10^4^ in the other panels), reducing mutational input. In this and the following two figures, mutations are turned off for the last 10^5^ generations, above the dashed line, showing the alleles maintained by balancing selection. Furthermore, the width is proportional to allele count, and colour indicates the sex-specific promoter affinities *α*_♀_ and *α*_♂_ according to the inset colour scale (where circles indicate evolutionary endpoints), and time (vertical axis) is shown on a log-scale, giving higher resolution for the faster initial evolution, and Roman numerals indicate the corresponding parameter combinations in Figures 6–7 (except for panel **iic**, which is not represented because it uses a different population size).

Simulations are run for 2 *×*10^5^ generations, followed by an additional 10^5^ generations without mutational input. We then count the remaining alleles to assess the persistence of polymorphism in the absence of recurrent mutation. The effects of mutation rate and population size are explored more systematically in Supplementary Figures S3–S5, using longer simulations of 10^6^ generations followed by 2*×* 10^5^ generations without mutational input.

To be able to investigate the effect of different constraints on sex-specific promoter evolution, we consider two alternative models. In the *shared-promoter model*, the promoter is characterized by a value *α*, but sex-specific transcriptional environments modulate its effect: 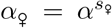 and 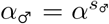, where *s*_♀_, *s*_♂_ *∈* [0, 1] are parameters describing the sex-specific sensitivities to the promoter affinity (e.g., via sex-biased transcription factors). This could, for example, be due to direct or indirect effects of major-effect transcription factors related to sex-determination, such as SRY in mammals, or *Doublesex* in insects. In this case, mutational effects on log(*α*) are drawn from a one-dimensional Gaussian distribution.

In the *independent-affinity model*, affinities in females and males, *α*_*i*, ♀_ and *α*_*i*, ♂_, evolve independently, reflecting sex-specific regulation of the same upstream region (e.g., sex-specific epigenetic regulation). In this case, mutations affect the vector (log *α*_♀_, log *α*_♂_) and, accordingly, their effects are drawn from a two-dimensional Gaussian distribution with correlation *ρ*.

In both cases, evolving *α*-values are restricted to the interval [10^*−*2^, 10^2^], and initial values are set to 1. Because affinities are strictly positive, we model mutations as proportional changes (i.e., additive steps in log(*α*)), drawing mutational effects from a normal distribution of log(*α*)-values with expected step size log(2) (so that *α*_min_ and *α*_max_ are ~13 mutational steps apart). For the upstream promoter sequence, we assume a somewhat higher effective per-gene mutation rate, namely *µ*_*α*_ = 5*×* 10^*−*5^, to reflect the additional mutational processes that affect regulatory regions, such as repeat-associated mutations, CpG enrichment and mutations associated with DNA methylation (Fan and Chu, 2007; Zhou et al., 2020; Kaiser et al., 2021; Liao et al., 2023).

## Results

Our results can be summarised in three points. First, in the absence of dominance evolution, we recover the established result that the conditions for stable biallelic polymorphism become increasingly restrictive under weak or asymmetric selection (e.g., Kidwell et al., 1977; Connallon and Clark, 2012; Arnqvist et al., 2014; Flintham et al., 2025), and that evolutionarily stable polymorphisms require strong and symmetric selection (Flintham et al., 2025). Such evolutionarily stable polymorphisms set the stage for the evolution of dominance (Fisher, 1931), and we show that, if sex-specific *cis*-promoter affinities are allowed to evolve, beneficial sex-specific dominance evolves and reduces segregation load. Second, coevolution of allelic values *x* and their sex-specific promoter affinities *α*_♀_ and *α*_♂_ can result in polymorphism also under weak or asymmetric selection. Importantly, we show that the coevolutionary feedback between allelic divergence and sex-specific dominance can be initiated in three ways: (i) during a selective sweep, (ii) during an evolutionarily transient protected polymorphism, or (iii) from genetic variation under mutation–selection– drift balance. Scenario (ii) is in line with existing theory (Spencer and Priest, 2016), whereas scenarios (i) and (iii) have previously been considered unlikely settings for the evolution of dominance (Haldane, 1956; Otto and Bourguet, 1999). Third, under weak or asymmetric selection, coevolution in *x* and *α* can lead to polyallelic polymorphism (PAP). In the following, we describe these results in detail.

We investigate the coevolution of allelic values *x* and their sex-specific promoter affinities *α*_♀_ and *α*_♂_, which together determine a phenotypic trait value *z* under sexually antagonistic selection. Selection on promoter affinities favours beneficial dominance (increasing *D*_♂_ or decreasing *D*_♀_) (Supplementary material S3.1). However, such selection requires polymorphism in *x*, since promoter affinities affect fitness only in individuals heterozygous for alleles differing in *x*.

Considering the evolution of allelic value *x*, we find that a single evolutionarily attracting value *x*\* exists. Starting from a monomorphic population far away from the evolutionary attractor, directional selection on the allelic value *x* causes allelic values to converge toward *x*\* through successive invasion and fixation of mutant alleles that are ever closer to *x*\*. The attractor *x*\* lies between the female and male optima (Eq. S26, see also Flintham et al. 2025), at the exact midpoint under symmetric selection and otherwise closer to the optimum of the sex under stronger selection (black dots in Fig. 1). The phenotypic value *z*\* = *x*\* maximises geometric mean fitness across the two sexes (Fig. 1). At *x*\*, the opposing directional selection gradients in the two sexes balance, with females favouring lower *x* and males favouring higher *x*. This holds for both autosomal loci (in line with previous work, e.g. Charnov, 1982; Leimar, 2001) and X-linked loci (Supplementary material S3.2.2).

Selection on promoter affinities is present whenever mutant and resident alleles coexist in heterozygotes. Far from *x*\*, however, directional selection on *x* is strong and such coexistence is generally restricted to short selective sweeps, leaving little opportunity for affinity evolution to affect the evolutionary dynamics. Consequently, changes in promoter affinity that arise during these early selective sweeps generally do not persist. As the population approaches *x*\*, directional selection weakens and mutant–resident coexistence can persist for longer, increasing the opportunity for selection on promoter affinities to become consequential. Importantly, this does not require an established balanced polymorphism: below we show that the coevolutionary feedback can be initiated during a selective sweep, during an evolutionarily transient protected polymorphism, or from mutation–selection–drift variation near *x*\*. We first consider weak or asymmetric selection before turning briefly to the less interesting case of strong and symmetric selection.

### Weak or asymmetric selection

Under this regime, *x*\* is a fitness maximum that would be reached and then be an evolutionary endpoint as long as alleles act additively (Fig. 3**i**). However, as the population evolves closer to the attractor *x*\*, coevolution in *x* and *α* can lead to the emergence of polymorphism in *x* associated with sex-specific dominance via the three scenarios listed above.

We start by describing scenario (ii). In the neighbourhood of *x*\*, there always exists a set of allelic values *x*_1_ and *x*_2_ with *x*_1_ *< x*\* *< x*_2_ that can coexist in a protected polymorphism due to heterozygotes having higher fitness when averaged over the two sexes (i.e., marginal overdominance; Supplementary materials S3.2.5 and S4; see also Sellis et al. 2011). This result provides a direct connection to the classical coexistence conditions for sexually antagonistic alleles (Kidwell et al., 1977). At *x*\*, the opposing female and male selection gradients are equal in magnitude, such that small allelic deviations on either side of *x*\* generate locally balanced sex-specific fitness effects. Gradual evolution toward *x*\* therefore naturally brings the population into the increasingly narrow region of parameter space permitting protected polymorphism under weak sexually antagonistic selection.

In the absence of beneficial sex-specific dominance, alleles closer to *x*\* can subsequently invade, so these polymorphisms are evolutionary transient as they are expected to eventually be replaced by *x*\*. However, this process is slow because selection is weak in the neighbourhood of *x*\*, resulting in long periods of coexistence, as illustrated in Fig. 3**i**. Importantly, during these transient episodes of polymorphism in *x*, selection favours mutations in the promoter affinities *α*_♀_ and *α*_♂_ that shift heterozygote phenotypes toward their sex-specific optima (Supplementary material S3.1). This route is illustrated in Figure 3**iib**: shortly after a protected polymorphism has formed around *x*\* (around generation 10^3^), the allele more beneficial in males evolves a higher affinity in males and a lower affinity in females (indicated by orange colouration), while the reverse occurs for the allele more beneficial in females (indicated by blue-green colouration).

The emergence of sex-specific dominance can have a significant effect on the direction of selection acting on the allelic values *x*, changing selection on coexisting lineages from convergent to divergent. This changes an evolutionarily transient polymorphism into an evolutionarily stable one. This can be seen in Figure 3**ii**–**iib** and in the top row of Figure 5, where the *x*-values of the two coexisting lineages diverge after sex-specific affinities have evolved. Specifically, any BAP in the neighbourhood of *x*\* experiences disruptive selection on the allelic values if the promoter affinities have evolved such that Condition S43 is fulfilled (Condition S45 for the X-linked case). Under symmetric selection strength (*σ*_♀_ = *σ* = *σ*_♂_) and *δ* = 1 this condition simplifies to

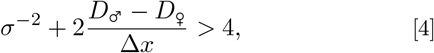

where the dominance parameters *D*_♂_ and *D*_♀_ (Eq. 2) express the direction and magnitude of dominance between two alleles in the two sexes, respectively, and Δ*x* = *x*_*i*_ *− x*_*j*_ *>* 0, where we index the alleles such that *x*_*i*_ *> x*_*j*_. A positive *D* indicates dominance of the allele with the larger allelic value, which is beneficial in males. Conversely, a negative *D* indicates dominance of the allele with the smaller allelic value, which is beneficial in females. Whenever *D*_♂_ ≠ *D*_♀_, dominance is sex-specific. Positive values of *D*_♂_ *− D*_♀_ mean that the higher-valued allele is more dominant in males than in females, and equivalently that the lower-valued allele is more dominant in females. We refer to this as beneficial sex-specific dominance. Importantly, dominance reversal is not required for Condition 4 to hold: divergence can also be favoured when dominance is in the same direction in both sexes, provided that it is stronger in the sex where it is beneficial (*D*_♂_ *> D*_♀_) (see Spencer and Priest, 2016, for a qualitatively similar conclusion from a two-allele, two-locus model).

There is another important insight one can glean from Condition 4. Since the left-hand side increases with the ratio (*D*_♂_ *− D*_♀_)*/*Δ*x*, only a small amount of beneficial sex-specific dominance is required for similar alleles (small Δ*x*) to experience disruptive selection. This generates a positive feedback loop: divergence in *x* increases the strength of selection for sex-specific dominance, while evolving sex-specific dominance promotes further divergence in *x*.

We now turn to scenario (i), in which sex-specific dominance evolves during a selective sweep. For selective sweeps far from *x*\*, with the resident and invading alleles positioned outside the interval between the sex-specific optima (open dots in Fig. 1), an invading allele *A* closer to *x*\* than the resident *a* has higher fitness in both the female and male contexts, such that *w*_*k*_(*a, a*) *< w*_*k*_(*A, a*) *< w*_*k*_(*A, A*) for *k ∈ {*♀,♂*}*. This corresponds to the classical setting in which dominance modifiers act only during the transient period of heterozygosity generated by the sweep, leaving little opportunity for promoter evolution (Haldane, 1956; Otto and Bourguet, 1999). Protected polymorphism between such alleles is not possible.

However, once selective sweeps involve pairs of alleles lying between the sex-specific optima, the situation is qualitatively different. Here, selection represents a tug-of-war between opposing sex-specific selection gradients: of two alleles differing in *x*, the allele with the lower value is more beneficial in females, while the allele with the higher value is more beneficial in males. Thus, an invading allele *A* positioned closer to *x*\* has higher homozygote fitness in one sex but lower homozygote fitness in the other, such that *w*_♂_ (*a, a*) *< w*_♂_ (*A, A*) and *w*_♀_ (*a, a*) *> w*_♀_ (*A, A*), or vice versa. Because each allele is favoured in a different sex, the net selective advantage of the invading allele is reduced, prolonging mutant–resident coexistence and thereby increasing the opportunity for promoter evolution. A *cis*-promoter mutation causing sufficient beneficial sex-specific dominance can then increase heterozygote fitness in the two sexes such that the alleles become mutually invadable, turning the selective sweep into a protected polymorphism.

Condition 4 describes this transition locally, for similar alleles in the neighbourhood of *x*\*. Under additivity, only alleles on opposite sides of *x*\* can coexist in a protected polymorphism, but when Condition 4 is fulfilled, mutual invadability extends locally to pairs of alleles that lie on the same side of *x*\* (compare Figure S7**B** with **A**). Thus, sufficiently beneficial promoter affinities can turn a selective sweep into a protected polymorphism before *x*\* is reached and initiate the feedback between sex-specific dominance and allelic divergence. This is illustrated under both weak and strong selection in Figures 3**ii** and 4**iv**. This route is more common under our baseline assumption that promoter affinities evolve more rapidly than allelic values (Supplementary Fig. S3) than when the mutation rate in *x* is higher (Supplementary Fig. S4). In the latter case, the feedback is more often initiated during a transient protected polymorphism around *x*\* (scenario ii).

Finally, in scenario (iii), the feedback begins from variation maintained around *x*\* by mutation–selection–drift balance. Here, the population first reaches the vicinity of *x*\* and remains effectively monomorphic, while low-frequency mutant alleles continue to arise. These mutants provide heterozygotes on which selection on promoter affinities can act. For small-effect mutants near *x*\*, a mutant displaced in either direction is favoured in one sex and disfavoured in the other. Beneficial sex-specific dominance can therefore reduce its phenotypic effect where detrimental while increasing it where beneficial. Once sufficiently strong, this changes local selection from convergent to divergent (Condition 4), causing the mutant and resident to become mutually invadable and initiating the same coevolutionary feedback as in scenarios (i) and (ii). This route was not observed under our baseline population size and mutation rates because diverging polymorphism emerged earlier from a selective sweep or transient protected polymorphism. It does occur under the smaller population size *N* = 5 *×* 10^3^, where reduced mutational input allows the population to reach *x*\* before the feedback is initiated, as illustrated in Figure 3**iic**. More generally, reducing population size increasingly delays and restricts allelic diversification, as lower mutational input causes the population to remain close to *x*\* for longer and stronger genetic drift reduces the efficacy of selection (Supplementary Fig. S5).

To investigate further how scenario (iii) can delay the onset of the evolutionary feedback, and how this depends on mutational input by varying *N*, Figure 5 compares simulations with the baseline population size (left column) to simulations with a smaller population size (right column), while keeping the mutation rate fixed. Affinity evolution is switched on only after 10^4^ generations. In the top row, a BAP has formed around *x*\* before affinity evolution begins, whereas in the bottom row the population is effectively monomorphic near *x*\*. From a preexisting BAP, sex-specific dominance evolves rapidly once affinity evolution is allowed, independently of *N*, and triggers divergence in *x*. When the population is effectively monomorphic, the feedback is delayed under smaller *N*, where lower mutational input generates fewer low-frequency variants and hence fewer heterozygotes on which selection on promoter affinities can act. At *N* = 5 *×* 10^4^, standing variation around *x*\* is instead sufficient for the feedback to begin with little delay. Thus, an established balanced polymorphism is not required for sex-specific dominance to evolve, but the amount of standing allelic variation strongly affects how rapidly the feedback is initiated.

Supplementary Figures S3–S5 investigate the robustness of these dynamics to mutational input. In Supplementary Figure S3, we reduce our baseline per-gene mutation rates by up to two orders of magnitude while maintaining *µ*_*α*_ = 10*µ*_*x*_. In Supplementary Figure S4, we repeat this analysis for the opposite relative mutation rates, *µ*_*x*_ = 10*µ*_*α*_. When *µ*_*α*_ *> µ*_*x*_, the coevolutionary feedback is more commonly initiated during a selective sweep (scenario i), whereas when *µ*_*x*_ *> µ*_*α*_ it is more commonly initiated during a protected polymorphism around *x*\* (scenario ii). Lowering mutational input slows down evolution and reduces the number of diversification events, but the coevolutionary feedback still occurs in most simulations even at the lowest mutation rates considered. At very low mutational input, the population more often first reaches the vicinity of *x*\*, such that the feedback is instead initiated from mutation–selection–drift variation (scenario iii). Reducing mutational input by decreasing population size has a qualitatively similar effect, while additionally increasing the importance of genetic drift (Supplementary Fig. S5). Allelic diversification becomes increasingly delayed and the number of diversification events decreases as *N* is reduced, resulting in lower final allelic diversity. At extremely small Δ*x*, however, selection itself becomes weak relative to drift.

Additionally, Supplementary Figure S6 investigates the effect of increasing mutational correlation *ρ* between female and male affinities. This has only a modest effect over most of the considered parameter range. The evolutionary dynamics slow considerably only as *ρ* approaches one, delaying the evolution of beneficial sex-specific dominance and subsequent allelic diversification.

### Strong and symmetric selection

Under sufficiently strong and symmetric selection, the attractor *x*\* is invadable by nearby mutant alleles (for the exact condition see Eq. S31, and Flintham et al. 2025 for an identical result). It is thus an evolutionary branching point in the sense of Geritz et al. (1998). Nearby mutant alleles can invade *x*\* and coexist with it in a BAP maintained by balancing selection. The two coexisting alleles then experience disruptive selection, leading to two divergent allelic lineages. In this case, diversification in *x* does not require evolution in the promoter affinities, as illustrated in Figure 4**iii**. Under additivity, divergence is driven by increasing homozygote fitnesses when averaged over the two sexes, while heterozygotes evolve lower fitness (Supplementary material S4). However, when promoter affinities are allowed to evolve, sex-specific dominance emerges readily and increases heterozygote fitness (Fig. 4**iv**). In the simulation shown, this coevolution begins already during the selective sweep toward *x*\*.

**Figure 4.**
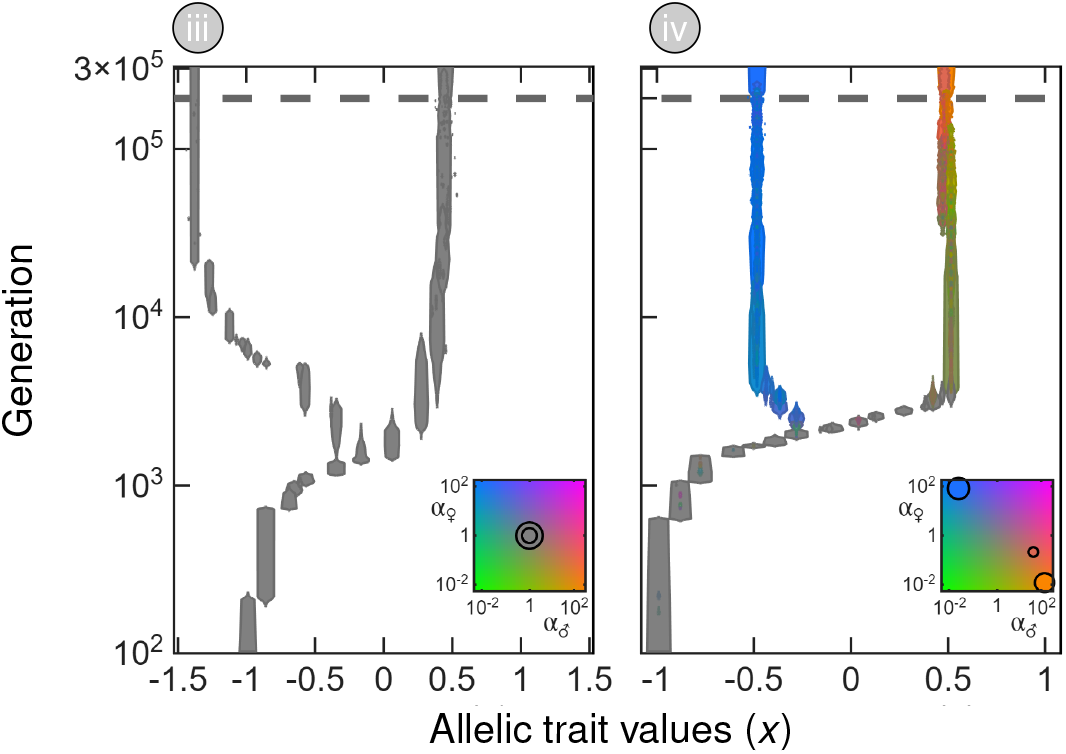
Individual-based simulations of allelic evolution at an autosomal locus under strong sexually antagonistic selection 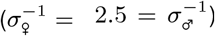. In panel **iii**, promoter affinities are fixed and alleles therefore act additively. The evolutionary attractor *x*\* = 0 is an evolutionary branching point, and the two lineages diverge without evolution of dominance; the resulting BAP is asymmetric, with one allele overshooting one sex-specific optimum (see Supplementary material S5). In panel **iv**, the sex-specific promoter affinities *α*_♀_ and *α*_♂_ can evolve. Beneficial sex-specific dominance emerges already during the selective sweep toward *x*\* and subsequently increases heterozygote fitness as the two allelic lineages diverge toward the sex-specific optima.

**Figure 5.**
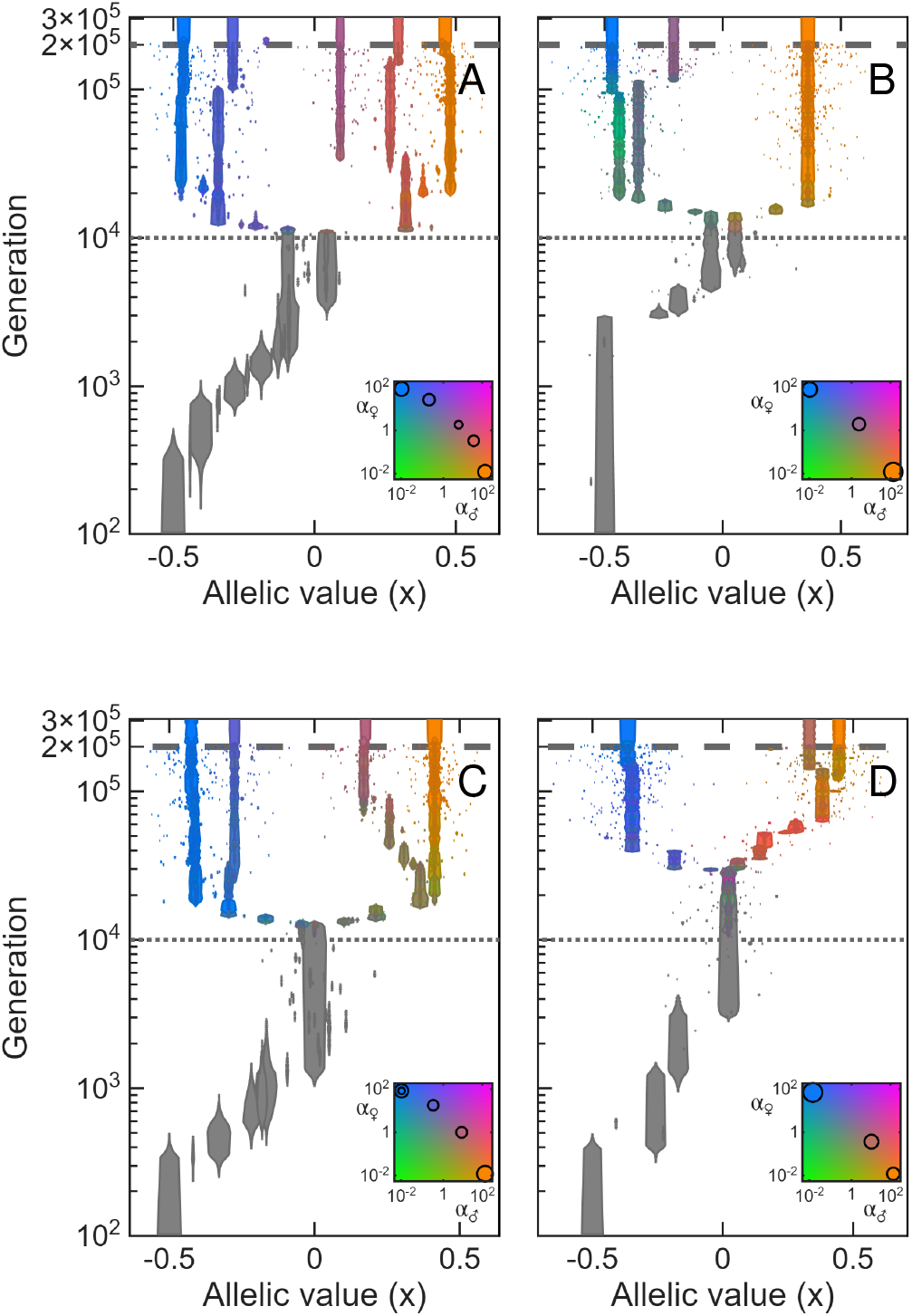
Individual-based simulations showing how transient protected polymorphism affects the emergence of sex-specific dominance. Simulations start with fixed promoter affinities, and affinity evolution is switched on at generation 10^4^ (dotted horizontal line). Unlike the other simulations, these are initialized 0.5 units from *x*\*. We consider an autosomal locus under moderate sexually antagonistic selection 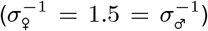, so the evolutionary attractor *x*\* = 0 is initiallya fitness maximum. In the top row (**A**–**B**), the population reaches a BAP consisting of alleles on opposite sides of *x*\* before affinity evolution begins, whereas in the bottom row (**C**–**D**) the population reaches a monomorphic state at *x*\*. The left column (**A, C**) uses the same population size as Fig. 3**iib.** The right column (**B, D**) uses a smaller population size (*N* = 5 *×*10^3^, as in Fig. 3**iic**) to reduce mutational input and emphasize the delay in diversification when starting from a monomorphic population at *x*\* (as in **D**).

### Evolution in polymorphic populations

We next examine how evolution proceeds after a diverging BAP has formed and first consider weak or asymmetric selection (Condition S31 not fulfilled). In this regime, continued evolution of promoter affinities can lead to repeated allelic diversification events in *x*. For example, Figure 3**iib** shows a simulation that ends with four coexisting alleles and evolved promoter affinities producing a dominance hierarchy (see inset), analogous to dominance hierarchies observed at naturally polymorphic loci (Billiard et al., 2021). Evolved sex-specific dominance shifts heterozygote phenotypes toward the corresponding sex-specific optimum and results in marginal overdominance: when averaged over the two sexes, heterozygotes have relatively high fitness, whereas homozygotes have relatively low fitness. As the number of coexisting alleles increases, the proportion of homozygotes diminishes, thereby reducing the segregation load due to low-fitness homozygotes. For instance, with four alleles at equal frequency, only 1*/*4 of individuals are low-fitness homozygotes (compared to 1*/*2 in a BAP), while the remaining genotypes are high-fitness heterozygotes. This pattern reflects the classical results of Kimura and Crow (1964), who showed that several alleles can be maintained when heterozygotes have similarly high fitness and homozygotes similarly low fitness.

In contrast, under strong and symmetric selection, generally a BAP consisting of the two optimal sex-specific alleles evolves, as shown in Figure 4**iv**. The inset in this figure shows that affinities coevolve such that each allele becomes dominant in the sex in which it is beneficial (dominance reversal; Fig. 2**C**). As a consequence, the heterozygote genotype expresses the sex-specific optimal phenotype in each sex and attains maximal fitness (white dots in Fig. 1). Due to the strong selection, intermediate allelic values *x* are strongly disfavoured because fitness drops sharply with increasing distance from the optima (Fig. 1**C**). Thus, heterozygotes carrying intermediate alleles have significantly lower marginal fitness than the heterozygote formed by the two specialist alleles, violating the near-equal heterozygote fitnesses that favour PAP (Kimura and Crow, 1964).

Figures 6 and 7 show the final number of coexisting alleles as a function of selection strength in males and females and under various constraints on the evolution of sex-specific affinities. In Figure 6 affinities are fixed so that alleles act additively. It shows that BAPs can emerge in most of the parameter space. However, from the lower panel, showing the maximum difference in allelic value among coexisting alleles, we can see that if Condition S31 is fulfilled (above and to the right of the solid white line), these BAPs correspond to diverged lineages (Fig. 4**iii**), while for weak or asymmetric selection (below and to the left of the solid white line), BAPs consist of similar alleles close to *x*\* (Fig. 3**i**) under stabilizing selection (Supplementary material S3.2.5). Simulations resulting in three or four coexisting alleles are due to transient dynamics between similar alleles.

**Figure 6.**
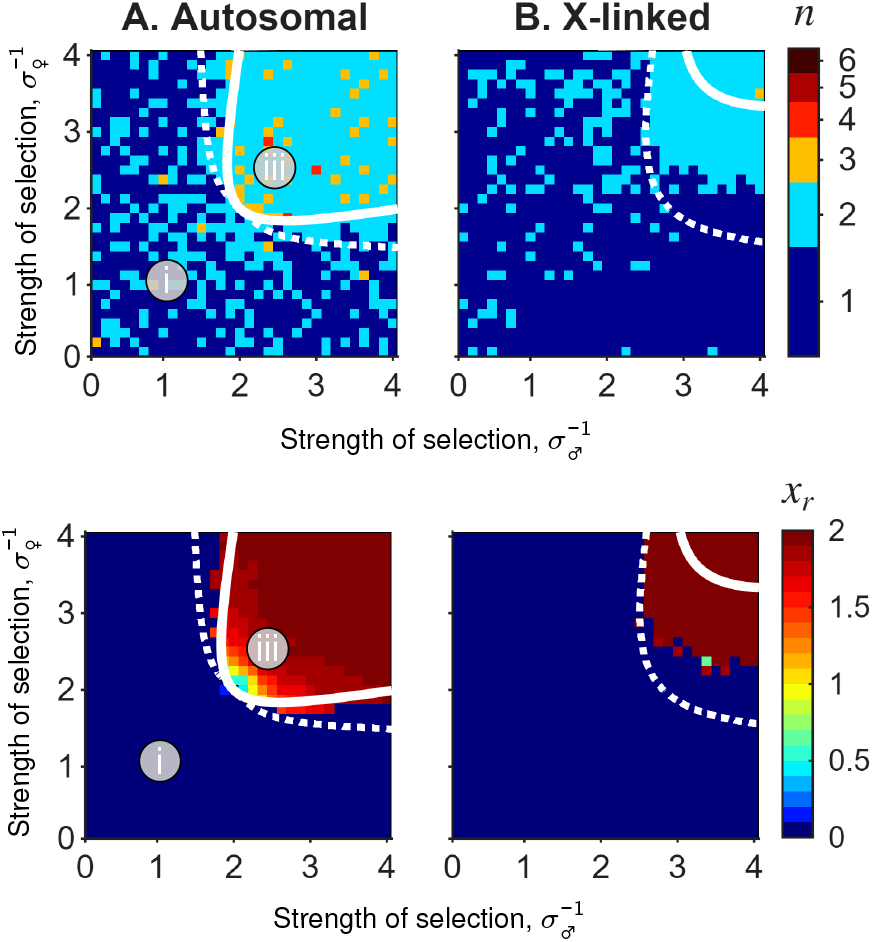
The number of coexisting alleles *n* (top row) and the maximum difference in allelic value *x* among coexisting alleles (bottom row) under sexually antagonistic selection without promoter evolution, shown as a function of the strength of selection in the two sexes 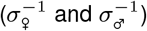 derived from individual-based simulations. **A** shows results for an autosomal locus and **B** for an X-linked locus. Solid white lines indicate the threshold for evolutionary branching under small mutational steps (Condition S31 for the autosomal locus and Condition S33 for the X-linked locus), whereas dotted white lines indicate the threshold at which *x*\* can be invaded by a large-effect mutant (Supplementary material S3.3). In this and the following figure, each pixel represents the outcome of a single simulation. Roman numerals mark parameter combinations for which the corresponding simulated evolutionary tree is shown in Figures 3 and 4.

**Figure 7.**
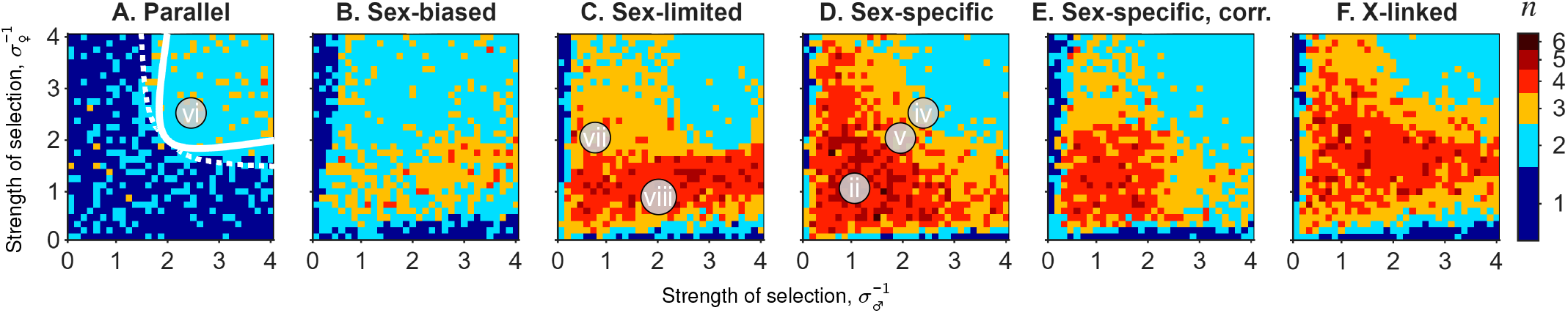
The number of coexisting alleles *n* under sexually antagonistic selection, shown as a function of the strength of selection in the two sexes 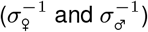 derived from individual-based simulations. Panels show results for different constraints on affinity evolution, as indicated above each panel. Panels **A**–**C** are based on the *shared-promoter model*. **A** assumes parallel dominance (*s*_♀_ = 1 and *s*_♂_ = 1, resulting in *α*_*i*♀_ = *α*_*i*, ♂_). **B** assumes female-biased dominance (*s*_♀_ = 1 and *s*_♂_ = 0.5, resulting in *α*_*i*, ♀_ = *α*_*i*_ and 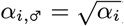. **C** assumes sex-limited dominance (*s*_♀_ = 1 and *s*_♂_ = 0, resulting in *α*_♀_ = *α*, while *α*_♂_ = 1 is insensitive to the promoter region). **D**–**E** are based on the *independent-affinity model*, where *α*_*i*, ♀_ and *α*_*i*, ♂_ evolve independently, with mutational correlation *ρ* = 0 in **D** and *ρ* = 0.9 in **E. F** shows results for an X-linked locus, where *α*_*i*, ♀_ evolves while males are hemizygous and carry a single allele *x*. The solid and dotted white lines are as in Figure 6A. Roman numerals mark parameter combinations for which the corresponding simulated evolutionary tree is shown in Figures 3, 4 and S1.

Figure 7 summarizes simulation outcomes under various constraints on affinity evolution. These simulations confirm that BAPs generically evolve under strong and symmetric selection, whereas weak or asymmetric selection can yield PAPs, with up to six coexisting alleles. Panels **A**–**C** use the *shared-promoter model* (see section with simulation details). In **A**, affinities are constrained to be identical across sexes (parallel dominance). Consistent with Condition S43, this does not promote polymorphism beyond the region expected under additivity. In **B** and **C**, promoter variation affects dominance fully in females, while males are increasingly less sensitive, generating increasingly sex-biased dominance. In **B**, dominance is sex-biased: male dominance follows the same ordering as in females but is weaker. In **C**, dominance is sex-limited to females, while alleles are additive in males. In both cases, polymorphism evolves across most of the parameter space, with PAP being more prevalent in **C** than in **B**. Panels **D** and **E** use the *independent-affinity model*. Panel **D** assumes no mutational covariance (*ρ* = 0), whereas in **E** female and male affinities are positively correlated (*ρ* = 0.9). Positive correlation slows the evolution of sex-specific dominance, but still produces a pattern of allelic polymorphism similar to the one without constraint. Supplementary Figure S6 shows that the effect of positive mutational correlations remains modest over a wide range of correlation values and becomes pronounced only as *ρ* approaches one.

#### X-linked loci

Comparing the two upper panels in Figure 6 shows that the condition for the emergence of diverging allelic polymorphism under additivity is more stringent for X-linked than for autosomal loci (see Eq. S33 for the exact condition, and Flintham et al. 2025 for a similar result). This mirrors earlier results from fixed two-allele models (Pamilo, 1979). The difference arises from the sex-biased inheritance of the X chromosome. A male-beneficial allele expressed in a surviving male is transmitted only to daughters, where it is detrimental, and can return to males only through those daughters. This weakens the effective selection favouring male-beneficial alleles. Female-beneficial alleles are less affected by this asymmetry, although their occurrence in female homozygotes also requires transmission through males, where they are detrimental. Overall, this inhibits allelic diversification and yields the more stringent evolutionary branching condition under additivity. However, because dominance is expressed only in females, allelic diversification is facilitated whenever *D*_♀_ *<* 0, corresponding to female-beneficial alleles being dominant in females. With evolving beneficial dominance, polymorphism therefore extends across a much broader range of selection strengths than under additivity, and the final number of alleles is similar to the autosomal case (Fig. 7**F**).

Finally, Conditions S31 and S33 are local conditions derived for mutations of small phenotypic effect, whereas mutations in our simulations have appreciable effects (the two sex-specific optima are approximately 13 average mutation steps apart). Larger mutational steps expand the parameter region in which polymorphism can arise through invasion of *x*\*, with the white dotted lines delimiting the region where *x*\* can be invaded by a large-effect mutant (see Supplementary material S3.3).

## Discussion

We show that sexually antagonistic selection can couple the evolution of sex-specific dominance to allelic diversification at a locus underlying a quantitative trait shared by the two sexes. Under additivity, our model recovers the established result that the conditions for stable biallelic polymorphism become increasingly restrictive under weak or asymmetric selection (e.g., Kidwell et al., 1977; Connallon and Clark, 2012; Arnqvist et al., 2014; Flintham et al., 2025). Allowing sex-specific promoter affinities to evolve changes this picture. Beneficial sex-specific dominance can initiate a coevolutionary feedback with allelic divergence, and this feedback can start during selective sweeps, during evolutionarily transient protected polymorphisms, or from variation maintained by mutation–selection–drift balance around the evolutionary attractor *x*\*. Under weak or asymmetric selection, repeated allelic diversification can ultimately generate polyallelic polymorphism (PAP).

Our results connect to classical theory on the evolution of dominance. Selection on dominance modifiers acts through heterozygotes (Fisher, 1931), and previous theory has therefore emphasized that dominance evolves particularly readily when balancing selection maintains heterozygotes for long periods (Otto and Bourguet, 1999). For sexually antagonistic loci, Spencer and Priest (2016) and Flintham (2025), similarly showed that a stable BAP can provide the heterozygosity needed for modifier evolution to produce beneficial sex-specific dominance. In their two-allele model, however, sex-specific fitness effects are fixed, so under weak selection stable polymorphism requires increasingly precise balance between the sexes (see also Kidwell et al., 1977). In our quantitative-trait model, gradual evolution toward *x*\* itself generates this balance and repeatedly creates transient protected polymorphisms on which selection on promoter affinities can act. Dominance evolution can then alter the evolutionary fate of the polymorphism that initially provides the heterozygotes, rather than merely evolving after balancing selection has already stabilized it: a transient polymorphism under stabilizing selection can turn into an evolutionarily stable polymorphism experiencing disruptive selection.

Importantly, an established protected polymorphism is not required before this feedback begins. In our simulations, it can start during selective sweeps once competing alleles lie between the sex-specific optima. This is in contrast to classical models of dominance evolution during selective sweeps, which typically consider selection around a single fitness optimum, where the invading allele is favoured over the resident (Haldane, 1956; Otto and Bourguet, 1999). Dominance can then alter heterozygote fitness but not the ultimately directional outcome of the sweep. The situation is qualitatively different under sexually antagonistic selection, where the alternative alleles in a sweep can be favoured in different sexes, creating a tug-of-war between opposing sex-specific selection gradients. Beneficial sex-specific dominance can therefore increase the phenotypic contribution of each allele where it is beneficial and reduce it where it is detrimental. If sufficiently strong, this can make the resident and invading alleles mutually invadable, turning an otherwise directional sweep into a protected polymorphism and extending a transient opportunity for dominance evolution into a sustained one.

The same principle allows the feedback to begin from mutation–selection–drift variation near *x*\*. Classical models consider mutations displaced from a single optimum, where they are deleterious in the same selective context and dominance can mainly mask their effects (Fisher, 1931; Otto and Bourguet, 1999). By contrast, individuals homozygous for *x*\* experience sexually antagonistic selection, and small-effect mutants are favoured in one sex and disfavoured in the other. Beneficial sex-specific dominance can therefore both reduce their phenotypic effect where detrimental and increase it where beneficial, allowing initially transient low-frequency variation to develop into a protected polymorphism and thereby create sustained selection on dominance.

Sex-specific dominance in trait expression thus significantly widens the conditions under which polymorphism evolves and is maintained. However, full beneficial dominance reversal—where alleles are fully dominant in one sex and fully recessive in the other in a manner aligned with selection—is not required. Around *x*\*, divergence can instead be promoted whenever beneficial dominance is sufficiently sex-specific, including when dominance remains in the same direction in both sexes but is stronger in the sex where it is beneficial (*D*_♂_ *> D*_♀_ in Condition 4), for example when allelic effects are additive in one sex but show some degree of dominance in the other (see also Brud, 2025). This is qualitatively consistent with Spencer and Priest (2016), who showed that a dominance modifier can invade when its benefit in one sex outweighs its cost in the other, and with Wittmann et al. (2017), who considered extremal rather than internal phenotypic optima but likewise found that relatively small differences in context-specific beneficial dominance can promote polymorphism under temporally fluctuating selection. Our model extends this principle by allowing both allelic values and dominance to evolve gradually, such that increasing beneficial sex-specific dominance can itself promote further allelic divergence.

Early theory showed that under additive genetic effects, the conditions favouring polymorphism are more restrictive for X-linked than autosomal loci (Pamilo, 1979), a result recovered in more recent models (Flintham et al., 2025). Classic theory nevertheless suggested that the X-chromosome could be enriched with SA loci (Rice, 1984), highlighting the importance of dominance for the evolutionary consequences of X-linkage. Subsequent models have shown that this expectation is not universal, particularly under sex-specific dominance reversal (Fry, 2010), and empirical evidence remains somewhat ambiguous (Dean and Mank, 2014; Ruzicka and Connallon, 2020). Our analyses recover the more restrictive conditions under additivity (compare Fig. 6**B** with **A**), but show that dominance can reverse this pattern, such that conditions for polymorphism become less restrictive in X-linked loci, particularly relative to autosomal loci under constrained dominance evolution (compare Fig. 7**F** with Fig. 7**B,C,E**). Importantly, dominance at an X-linked locus need not itself be sex-specific: because males are hemizygous, dominance expressed in females alone can generate the required asymmetry. Since such dominance can be at least as effective in promoting PAP as beneficial sex-specific dominance in autosomal loci (compare Fig. 7**F** with Fig. 7**B**–**E**), the X-chromosome may be enriched with polymorphic SA loci for this reason alone. Much previous theory on sexually antagonistic selection has considered dominance at the level of fitness, which can emerge from curved fitness functions even under additive allelic effects on the underlying phenotype (Connallon and Clark, 2014; Flintham et al., 2025). In our model, we additionally consider dominance at the level of trait expression, arising through allele-specific promoter affinities that can evolve to differ between the two sexes (see also Wittmann et al., 2017; Flintham, 2025). The Gaussian mapping from phenotype to fitness remains nonlinear and therefore also contributes to the resulting dominance at the fitness level. This distinction is relevant because, although dominance reversal for fitness can emerge from curved fitness landscapes (Fry, 2010; Manna et al., 2011; Connallon and Chenoweth, 2019), many well-documented cases of sex-specific dominance occur already at the phenotypic level. Examples include age at maturation in salmon (Barson et al., 2015), morphology in water striders (Fairbairn et al., 2023), immune function in fruit flies (Geeta Arun et al., 2021), migratory behaviour in rainbow trout (Pearse et al., 2019), and transcript abundance in fruit flies (Wayne et al., 2007; Puixeu et al., 2023; Mishra et al., 2022) and beetles (Kaufmann et al., 2024). Sex-specific dominance observed for fitness components (Grieshop and Arnqvist, 2018; Mérot et al., 2020) may therefore often emerge from underlying differences in dominance for trait expression, in line with our focus on the evolution of the genotype-to-phenotype map.

Sex-specific dominance is one of several mechanisms by which the genotype-to-phenotype map can become sex-specific and thereby reduce sexually antagonistic conflict. Sex-specific allelic effects, sex-limited expression, gene duplication, genomic imprinting and other forms of sex-specific regulation can similarly decouple male and female phenotypes (Bonduriansky, 2007). For instance, in a model similar to ours presented by Flintham et al. (2026), mutations can directly generate different allelic effects in females and males. Repeated mutation can therefore produce a single allele whose effect approaches the female optimum in females and the male optimum in males. Needless to say, such an allele will resolve conflict and thus displace polymorphism maintained by sex-specific dominance. Their model thus assumes that sex specificity in the allelic effect itself is directly evolvable, rather than modelling the mechanism by which such sex specificity arises. Here, by contrast, the underlying allelic value *x* remains shared between the sexes, and sex specificity in the genotype-to-phenotype map evolves through an explicit regulatory mechanism, namely sex-specific promoter affinities. Our model therefore asks how far sex-specific dominance can resolve the conflict when the underlying allelic effect remains constrained between the sexes.

This distinction also matters when moving from a single-locus to a polygenic architecture. Previous theory shows that genetic variation arising from disruptive selection tends in the long term to be concentrated in a small number of loci (Kopp and Hermisson, 2006; Van Doorn and Dieckmann, 2006; Yeaman, 2022). However, this expectation changes when sex-specific dominance is allowed to evolve. In additional multilocus simulations (Siljestam et al., 2026), we show that polymorphism is instead maintained across many loci, with allelic effects evolving toward an even distribution across the polygenic architecture. Thus, the coevolutionary feedback between sex-specific dominance and allelic diversification does not depend on a single-locus architecture. Across multiple loci, this feedback is iterated, with multiple polymorphic loci collectively generating stable sexual dimorphism and further reducing segregation load.

Unlike previous theory on sex-specific dominance, our model is not restricted to BAP. Under weak or asymmetric selection, repeated diversification can generate PAP, which further reduces segregation load because the fraction of low-fitness homozygotes declines as the number of coexisting alleles increases (Rice and Chippindale, 2002). Marginal overdominance is well known to promote the evolution and maintenance of PAP when heterozygotes have similarly high fitness (e.g., Kimura and Crow, 1964; Siljestam and Rueffler, 2024). In our model, under weak selection, fitness declines slowly with deviations from the sex-specific optima, allowing heterozygotes formed by a broad range of allelic values to attain similarly high fitness and thereby permitting multiple alleles to coexist. Under strong selection, by contrast, heterozygotes formed by alleles closest to the two sex-specific optima outcompete those involving intermediate alleles, favouring BAP. PAP has also been predicted under interlocus sexual conflict through a different mechanism (Gavrilets and Waxman, 2002; Haygood, 2004). The regulatory mechanism modelled here points particularly to regions affecting gene expression. Pronounced sexual dimorphism in gene expression is widespread in gonochorists (Ellegren and Parsch, 2007), and both theory (Connallon and Knowles, 2005; Rowe et al., 2018; Tosto et al., 2023) and empirical data (Innocenti and Morrow, 2010; Hollis et al., 2014; Wong and Holman, 2023) suggest that regulatory regions are often under, or have experienced a history of, SA selection. This could be true for regions affecting both sex-specific transcription and sex-specific splicing (Telonis-Scott et al., 2009; Rogers et al., 2021; Singh and Agrawal, 2023). Moreover, several studies have documented widespread sex-specific dominance in gene expression (Wayne et al., 2007; Meiklejohn et al., 2014; Puixeu et al., 2023; Mishra et al., 2022; Kaufmann et al., 2024), suggesting that regulatory regions may be enriched for loci showing segregating SA variation. Although a few specific cases of PAP have been documented in candidate SA genes (Barson et al., 2015; Schmidt et al., 2010; Hawkes et al., 2016; Wagstaff and Begun, 2005; Lonn et al., 2017), relevant empirical data are limited. However, several genomic regions with putative regulatory functions, such as mini- and microsatellites, show pronounced PAP and sometimes appear to be under balancing SA selection (e.g., Lonn et al., 2017). When superimposed on our results, these observations suggest that a sizeable fraction of standing SA genetic variance for fitness (Connallon and Matthews, 2019) may actually be caused by PAP in *cis*-regulatory regions (Wright et al., 2018; Kaufmann et al., 2024).

In summary, our results broaden the conditions under which sex-specific dominance can evolve and maintain sexually antagonistic variation. The coevolutionary feedback by which this happens need not begin from an established balanced polymorphism: it can start during selective sweeps, from evolutionarily transient protected polymorphisms, or from variation maintained by mutation–selection–drift balance. Once initiated, the feedback between promoter evolution and allelic diversification can repeat and ultimately generate PAP, particularly under weak selection. We suggest that time will thus sift novel SA mutations, preferentially retaining those that show beneficial sex-specific dominance either from the outset or acquire it through subsequent evolution. As a result, standing SA genetic variation for fitness, which is often sizeable (Connallon and Matthews, 2019), should be enriched for loci showing sex-specific dominance and PAP.

## Code availability

The MATLAB code for the individual-based simulations is publicly available on Zenodo at https://doi.org/10.5281/zenodo.23598283.

## Author contributions

Conceptualization: M.S., G.A. and C.R.; Methodology: M.S. and C.R.; Formal analysis: M.S.; Visualization: M.S.; Writing: M.S., G.A. and C.R.; Supervision: G.A. and C.R.; Funding Acquisition: C.R. and M.S.

## Supplementary Material to

### S1. Supplementary figures

**Figure S1.**
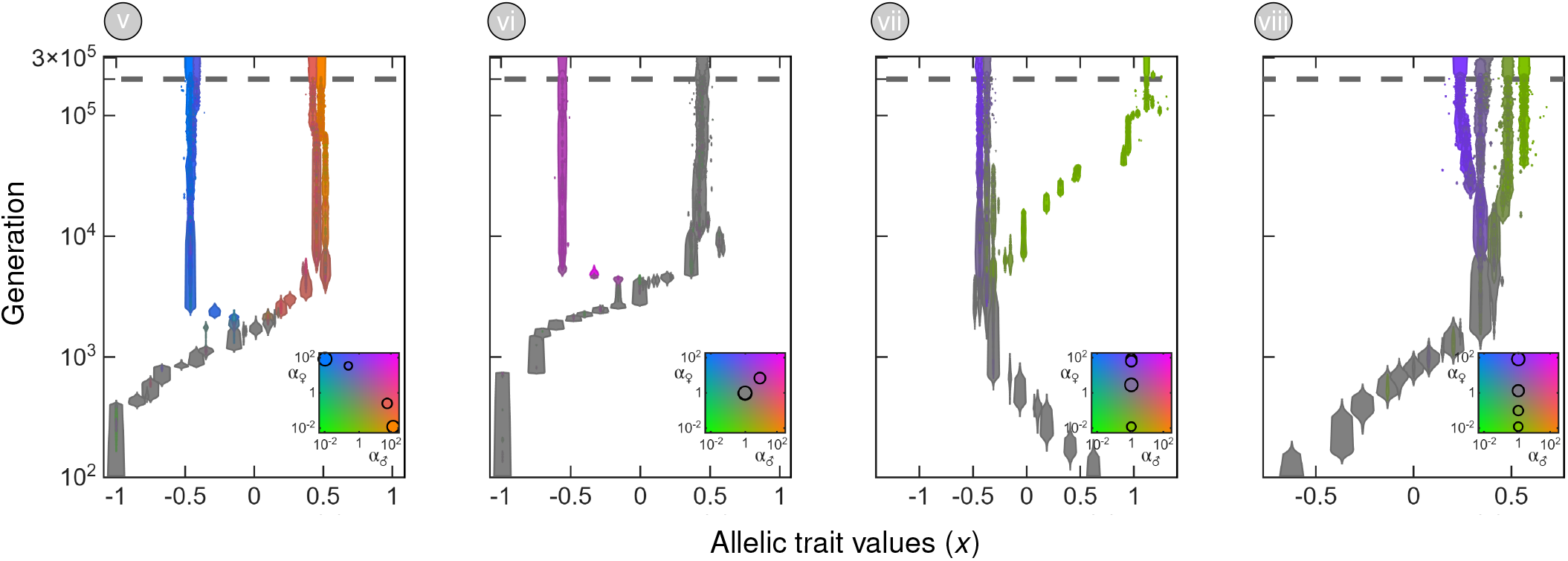
Individual-based simulations showing the evolution of allelic values at an autosomal locus under sexually antagonistic selection. Parameter values for each simulation are given by the location of the Roman numerals in Figure 7. Panel **v**: under the independent-affinity model and intermediately strong selection 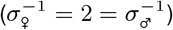, two allelic clusters evolve, each centred around one of the sex-specific optima. Panels **vi**–**viii** show evolution under the shared-promoter model. Panel **vi**: under parallel dominance (α_*i*,♀_ = α_*i*,♂_) and strong symmetric selection 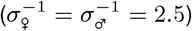, evolutionary branching produces a BAP, as predicted by the branching condition in the absence of sex-specific dominance (Condition S31). Panels **vii** and **viii** show evolution under modifier evolution limited to females for weak and asymmetric selection, with 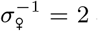 and 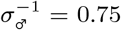 in panel **vii**, and 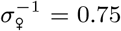 and 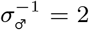 in panel **viii.** In panel **vii**, an allelic cluster around the female optimum co-occurs with one allele that overshootsthe male optimum. In panel **viii**, a PAP evolves that is tightly clustered around the male optimum.

**Figure S2.**
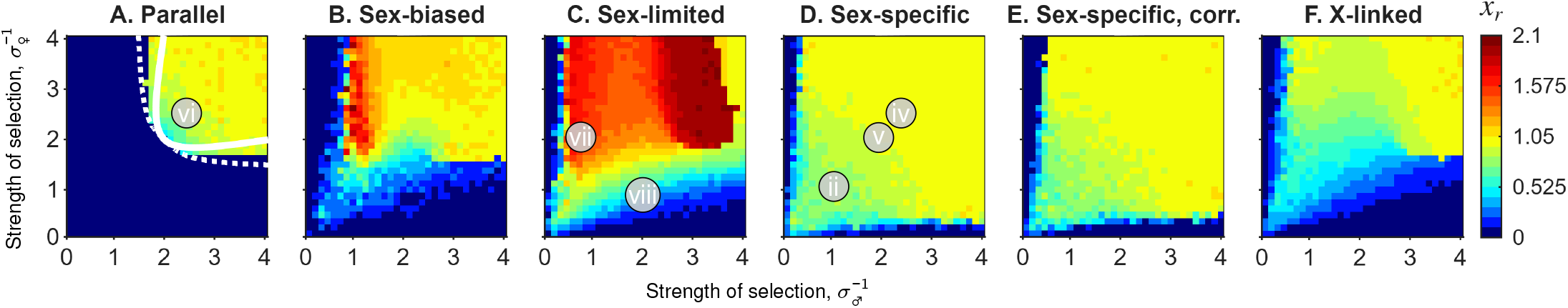
Maximum difference in allelic value *x* among coexisting alleles under sexually antagonistic selection, shown as a function of the strength of selection in the two sexes 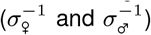 and derived from individual-based simulations. Each panel complements the corresponding panel in Figure 7. Roman numerals mark parameter combinations for which the corresponding simulated evolutionary trajectory is shown in Figures 3, 4, and S1.

**Figure S3.**
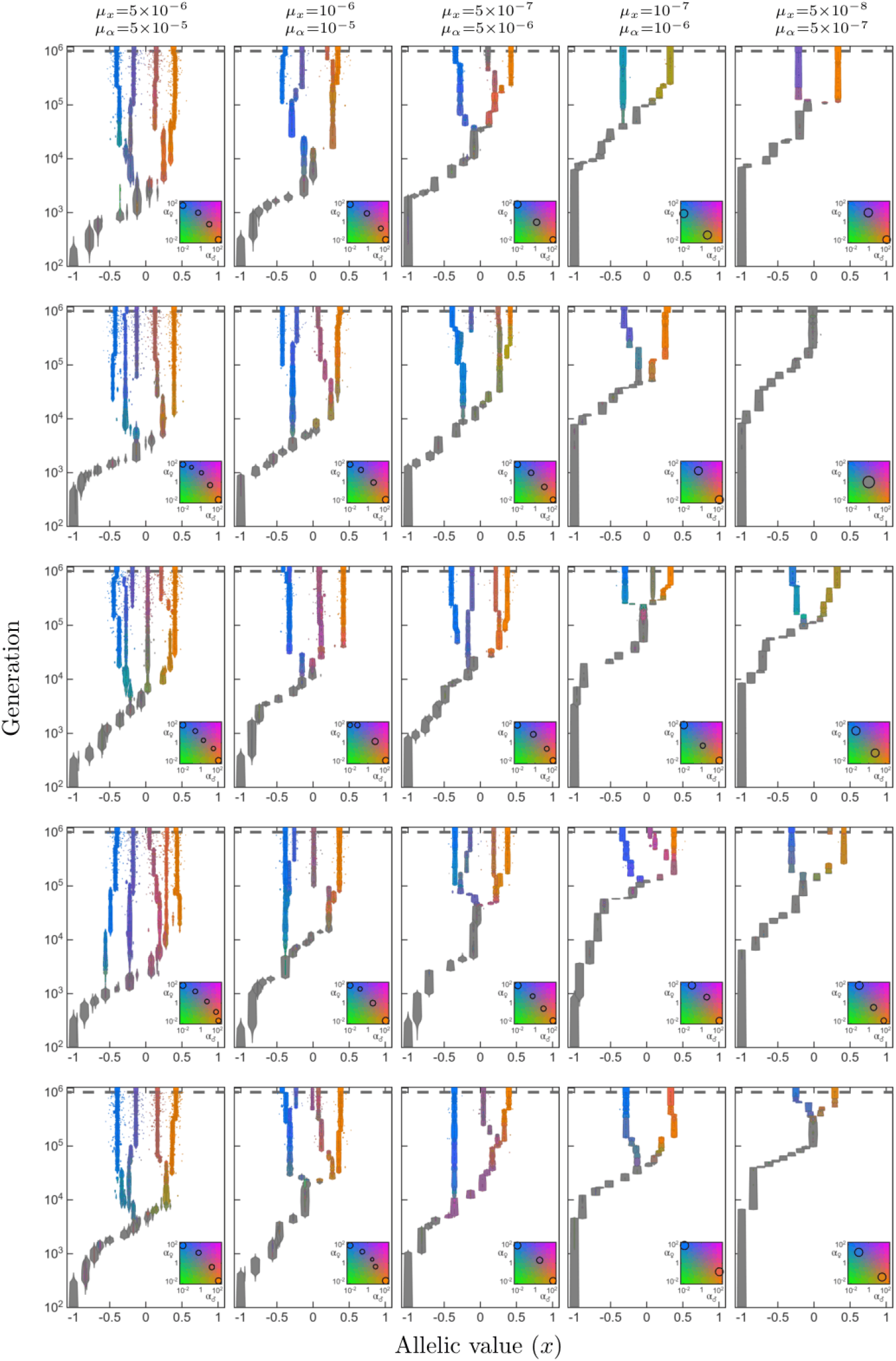
Evolutionary trajectories under decreasing mutational input. Each panel shows an individual-based simulation of the joint evolution of allelic value *x* and sex-specific promoter affinities under sexually antagonistic selection. Columns differ in per-gene mutation rate, with *µ*_*α*_ = 10 *µ*_*x*_, from the highest mutation rate on the left to the lowest on the right. Rows show replicate simulations using different random seeds. Simulations were run for 10^6^ generations with mutation, followed by an additional 2*×* 10^5^ generations without mutation. Across most mutation regimes, the coevolutionary feedback between beneficial sex-specific dominance and genetic polymorphism leads to allelic diversification. The exception is the second row at the lowest mutational input (*µ*_*x*_ = 5*×* 10^*−*8^, corresponding to approximately one mutation affecting *x* every 200 generations), where allelic values do not diverge from the attractor within the simulated time. The number of allelic diversification events decreases substantially at the lower mutation rates in the two right-most columns. Because promoter affinities mutate ten times more frequently than allelic values, the coevolutionary feedback is often initiated during selective sweeps before the population reaches *x*\*. At very low mutational input, coevolution can be initiated from mutation–selection–drift variation around a single allele close to *x*\*, resulting in an increasing delay before divergence starts.

**Figure S4.**
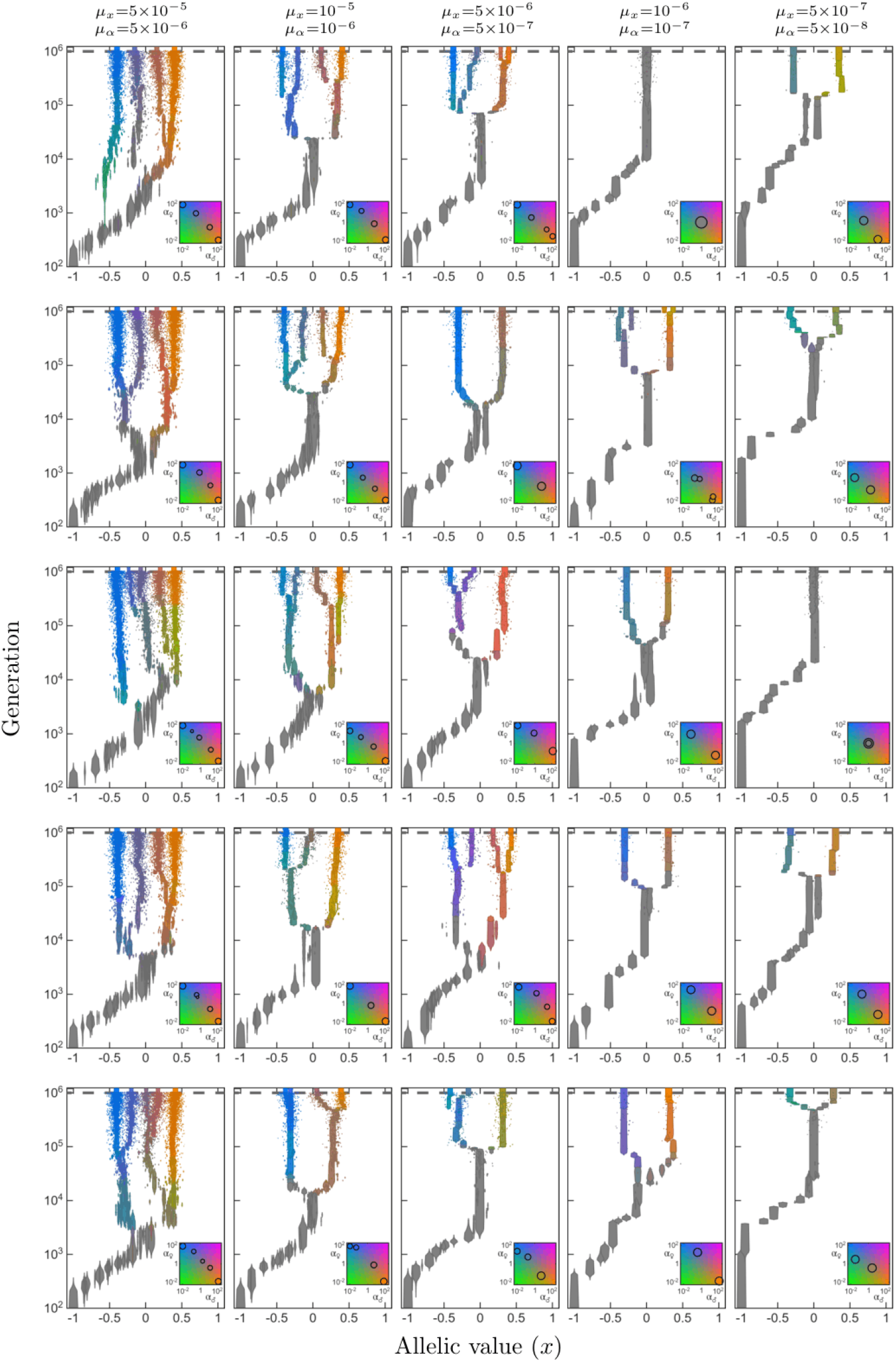
Evolutionary trajectories under decreasing mutational input, with the mutation rate of allelic values ten times higher than that of promoter affinities (*µ*_*x*_ = 10 *µ*_*α*_). Otherwise, simulations are as in Figure S3. Because allelic values mutate ten times more frequently than promoter affinities, sex-specific dominance more often evolves after transient protected polymorphisms have formed between alleles on opposite sides of *x*\*. At very low mutational input, coevolution instead initiates from mutation–selection–drift variation around a single allele close to *x*\*, resulting in an increasing delay before divergence starts. At each of the two lowest promoter-affinity mutation rates, corresponding to approximately one promoter mutation every 100 and 200 generations, respectively, one replicate fails to initiate the coevolutionary feedback within the simulated time.

**Figure S5.**
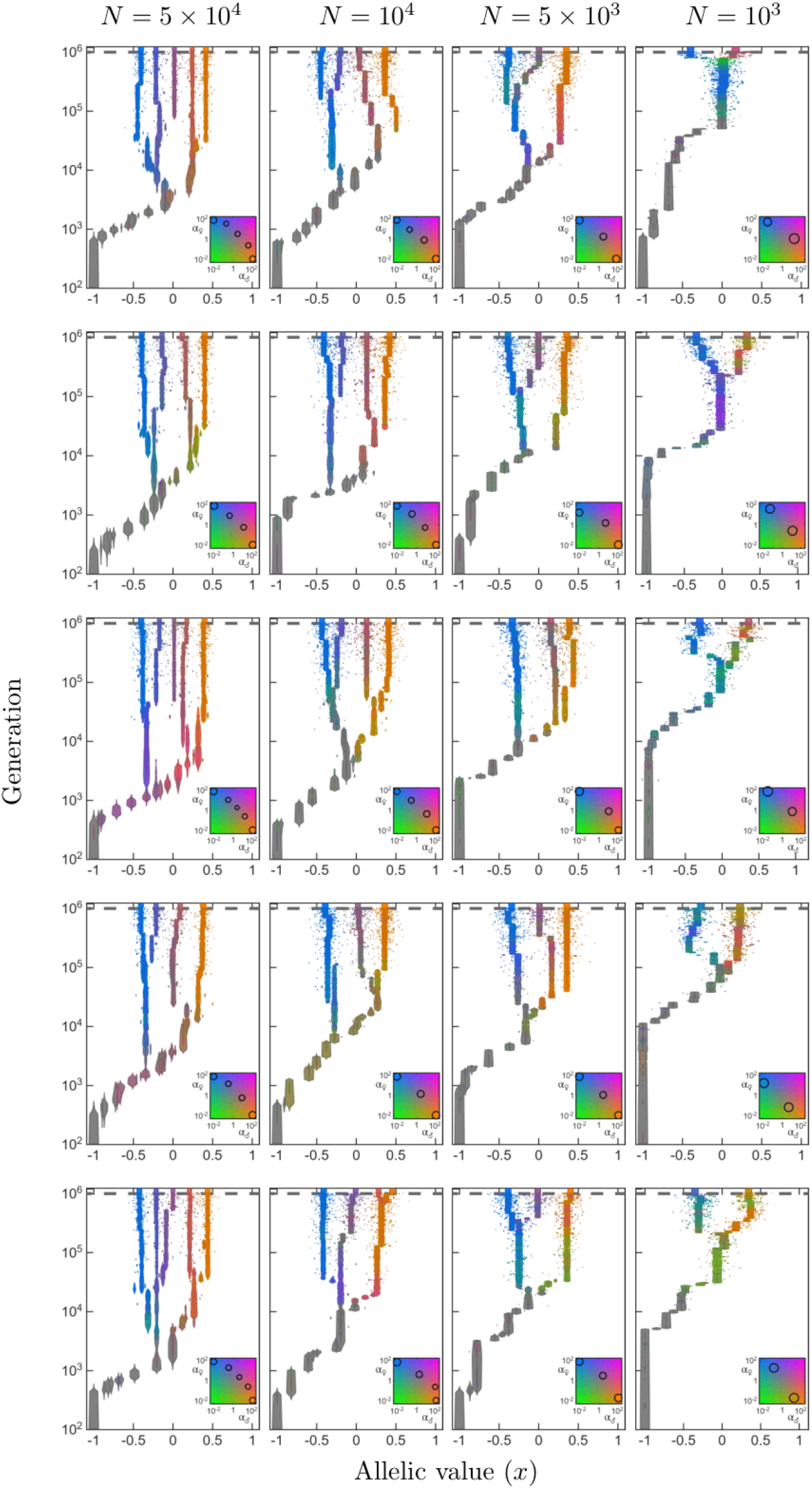
Evolutionary trajectories under decreasing population size. Columns differ in population size *N*, while rows show replicate simulations using different random seeds. Mutation rates are held constant at the baseline values *µ*_*x*_ = 5 *×*10^*−*6^ and *µ*_*α*_ = 5 *×*10^*−*5^. At larger population sizes, the coevolutionary feedback is initiated during selective sweeps or from transient protected polymorphisms around *x*\*, as in the baseline simulations. As *N* decreases, reduced mutational input makes the population increasingly likely to reach the vicinity of *x*\* before the feedback is initiated, such that coevolution instead begins from mutation–selection–drift variation around a single allele near *x*\*. Consequently, allelic diversification becomes increasingly delayed and the number of diversification events decreases, resulting in lower final allelic diversity. At the smallest population size, a new *x* mutation arises on average only once every approximately 100 generations. The increasing importance of genetic drift is also evident from promoter-affinity mutations drifting to fixation while the population is monomorphic in *x*, when variation in promoter affinity is selectively neutral.

**Figure S6.**
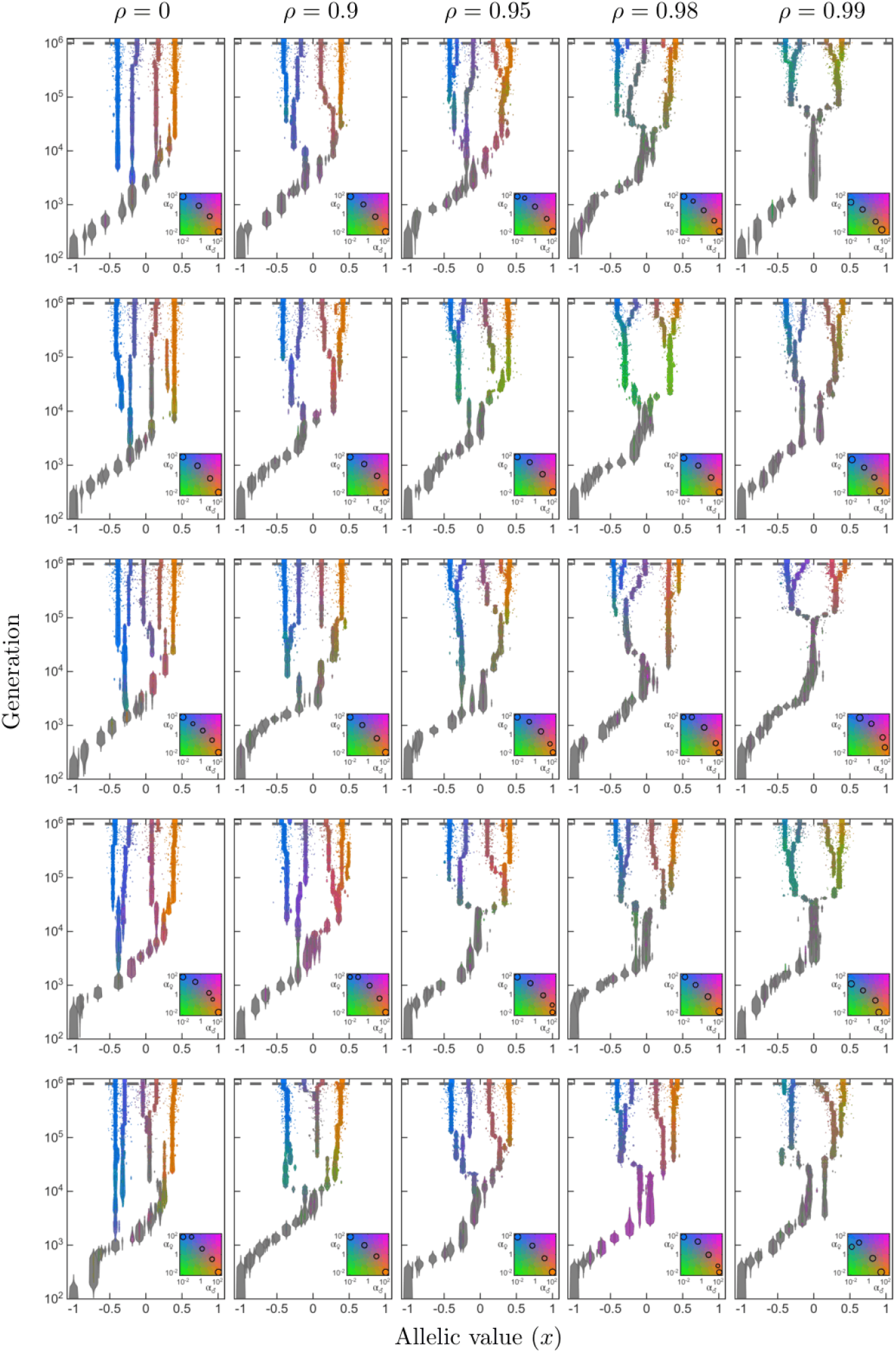
Evolutionary trajectories under increasing mutational correlation *ρ* between mutational effects on female affinities *α*_♀_ and male affinities *α*_♂_. Columns differ in *ρ*, while rows show replicate simulations using different random seeds. Mutation rates are held constant at the baseline values *µ*_*x*_ = 5 *×*10^*−*6^ and *µ*_*α*_ = 5 *×*10^*−*5^. Increasing mutational correlation generally has only a modest effect on the evolutionary dynamics until *ρ* approaches one. At very high correlations, mutations increasingly alter female and male affinities in parallel, slowing the evolution of beneficial sex-specific differences in dominance and thus delaying onset of the coevolutionary feedback and subsequent allelic diversification.

**Figure S7.**
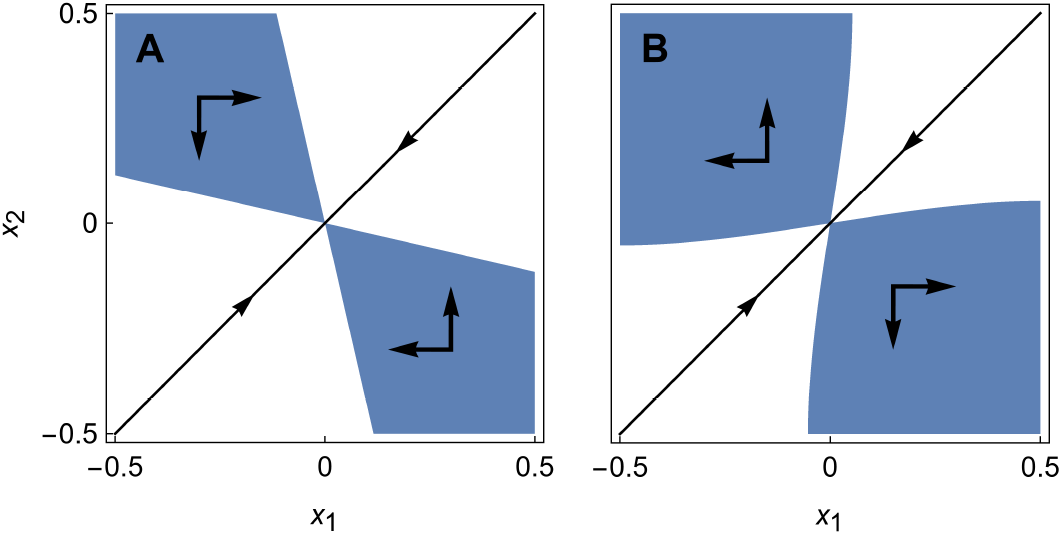
Coexistence near the singular allele *x*\* = 0. All combinations of two allelic values *x*_1_ and *x*_2_ that can mutually invade each other and therefore coexist in a protexted polymorphism are shown in blue. We prove in Section S3.2.5 that there always exists a set of coexisting alleles on opposite sides of *x*\*. In **A**, the condition for evolutionary branching is not fulfilled. As a consequence, coexisting alleles experience stabilizing selection, indicated by arrows pointing toward *x*\* = 0. In **B**, the condition for evolutionary branching is fulfilled, resulting in divergent selection acting on two coexisting alleles. This is indicated by arrows pointing away from *x*\* = 0. Alternatively, this figure can illustrate the effect of beneficial sex-specific dominance on the evolutionary dynamics of a BAP. Using the parameters from Figure 6**i** with weak selection 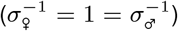, panel **A** corresponds to the absence of sex-specific dominance (*d*_♀_ = 0 = *d*_♂_). In this case, BAPs close to *x*\* are evolutionary transient as they experience stabilizing selection toward *x*\*. In contrast, panel **B** shows that with beneficial sex-specific dominance (*d*_♀_ = −2.5, *d*_♂_ = 2.5 as in Eq. S39; Condition S43 fulfilled), BAPs instead experiences disruptive selection away from *x*\*. See Geritz et al. (1998) for how to construct such coexistence plots.

**Figure S8.**
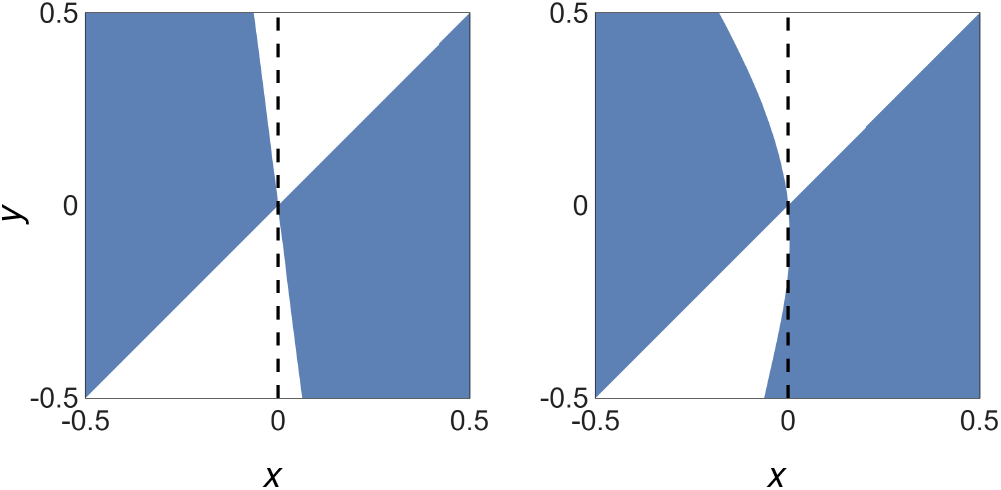
Pairwise invasibility plots (Geritz et al., 1998) illustrating local and global invadability of the singular allele *x*\* in the absence of dominance. Resident allelic values *x* are along the x-axis while mutant allelic values *y* are along the y-axis. **A.** Autosomal locus under moderate sexually antagonistic selection with 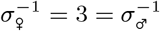, where *x*\* is globally uninvadable. **B**. X-linked locus under strong sexually antagonistic selection with 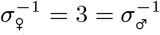, where nearby mutants cannot invade a resident at *x*\* but mutants with sufficiently lower allelic value can invade. Thus, a local evolutionary endpoint under small mutational steps need not be globally uninvadable.

**Figure S9.**
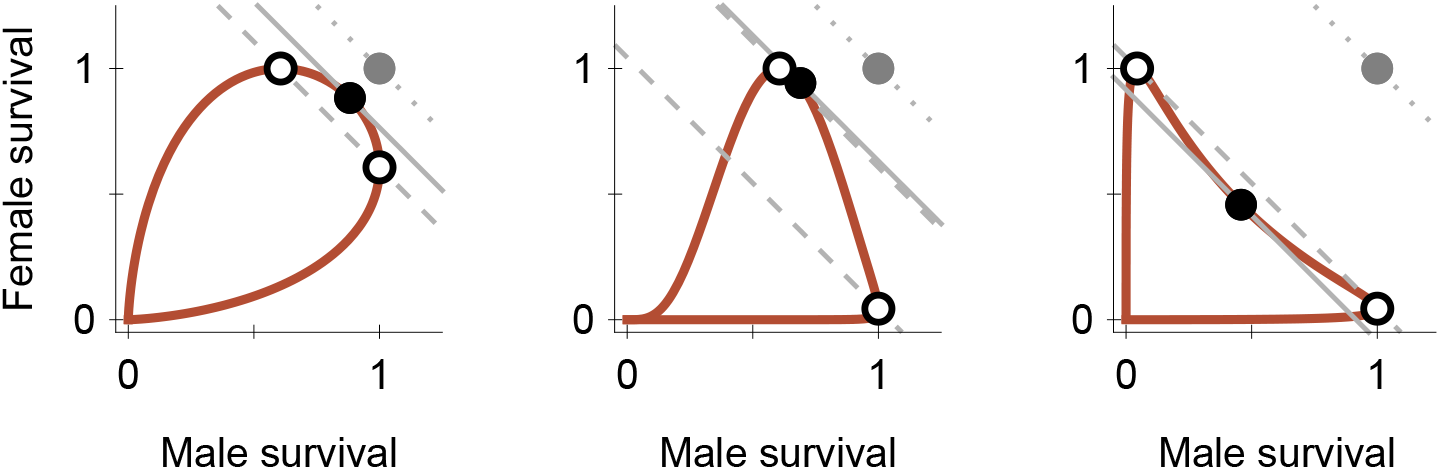
Sex-specific fitness (viability) *w* as a function of the phenotypic trait *z*, illustrating the same Gaussian trade-offs as in Figure 1, but with the two fitnesses shown as a parametric curve in *z* (cf. Levins, 1968, p. 16). Open dots indicate the phenotypic optima for females and males, while filled dots indicate the trait value that maximizes geometric mean viability across the two sexes. Contour lines of equal fitness averaged across both sexes (marginal fitness) are added in grey. Hatched lines indicate all combinations of female and male fitness that have the same marginal fitness as the phenotype that maximizes fitness in one sex. Solid lines do the same, but use the trait value maximizing geometric mean fitness as the reference. Finally, dotted lines pass through a filled grey dot corresponding to the female and male fitnesses of individuals heterozygous for the two specialist alleles under full beneficial dominance. Because females and males express different trait values under sex-specific dominance, these dots no longer lie on the parametric curve. In **A**, both sexes are under weak selection 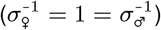, while in **B** females are under strong selection and males under weak selection 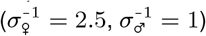. In **C**, both sexes are under strong selection 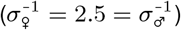.

#### S2. Mathematical analysis

We investigate whether selection at a locus subject to sexually antagonistic selection favours a single allele or leads to polymorphism, and how this is affected if these alleles can evolve sex-specific affinities at their corresponding promoter regions. Our study encompasses both autosomal and X-linked loci. We address this question through evolutionary invasion analysis using the adaptive dynamics framework (Metz et al., 1992; Geritz et al., 1998; Doebeli, 2011).

##### S2.1. Genetic dominance

In our model, selection acts on a trait that affects fitness through viability. Specifically, an individual’s fitness (viability) *w*_*k*_, for sex *k* ∈ {♀, ♂}, is determined by its phenotype *z*_*k*_. Each allele *i* is characterized by an allelic value *x*_*i*_ and a sex-specific promoter affinity *α*_*i,k*_ *>* 0. Because affinities are strictly positive and mutations are assumed to act proportionally on affinity, we work in the Supplementary Information with the corresponding log-affinities,

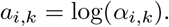

Multiplicative mutational effects on *α*_*i,k*_ then correspond to additive mutational effects on *a*_*i,k*_ (in the simulations, we use an expected mutational step size of log 2, corresponding to approximately a two-fold change in affinity *α* per mutation).

For a diploid individual carrying alleles *i* and *j*, the phenotype expressed in sex *k* is the affinity-weighted mean of their allelic values,

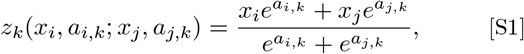

(Van Dooren, 1999). Here, 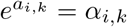 denotes the promoter affinity of allele *i* in sex *k*. When comparing two alleles, we index them such that *x*_*i*_ *> x*_*j*_ and define

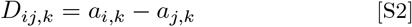

as their difference in log-affinity in sex *k*. Equation S1 can then be written more compactly as

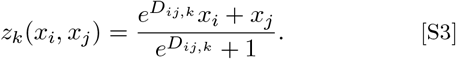

The dominance coefficient of the allele with the larger allelic value is therefore given by 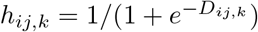. With this indexing convention, *D*_*ij,k*_ *>* 0 means that the allele with the larger allelic value has the larger promoter affinity and is phenotypically dominant, whereas *D*_*ij,k*_ *<* 0 means that the allele with the smaller allelic value is dominant. Hence, sex-specific dominance, *h*_*ij*,♀_ ≠ *h*_*ij*,♂_, follows if the relative size of the affinities differs between the sexes: *D*_*ij*,♀_ ≠ *D*_*ij*,♂_ (or, equivalently, 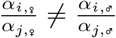. Note that for a homozygote, the phenotype function becomes independent of the affinities, and *z*_*k*_(*x, x*) = *x*.

For an X-linked locus, males are hemizygous, possessing only one allele inherited from their mother. For this case, we assume that the relationship between allelic value and phenotypic trait is also direct, *z*(*x*) = *x*.

### S2.2. Fitness

Fitness (or viability) is modelled as a Gaussian function of the difference between an individual’s phenotype *z*_*k*_ and the sex-specific optimal phenotype *θ*_*k*_,

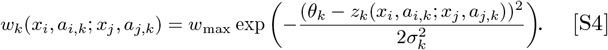

Here, *σ*_*k*_ determines the width of the Gaussian curve for sex *k*, and *w*_max_ denotes the fitness reached by an optimal phenotype. For hemizygous males, the viability function simplifies to

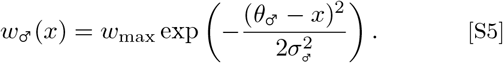

Note that *w*_max_ is omitted from all subsequent calculations, as it cancels out in both Equation S12 and Equation S13. Without loss of generality, we define the female and male optima as *θ*_♀_ = −*δ/*2 and *θ*_♂_ = *δ/*2, respectively, where *δ* gives the distance between these optima. For two alleles with *x*_*i*_ *> x*_*j*_, sex-specific dominance is beneficial when *D*_*ij*,♂_ *> D*_*ij*,♀_, such that the higher-valued allele is more dominant in males than in females, and equivalently the lower-valued allele is more dominant in females than in males. The special case *D*_*ij*,♀_ *<* 0 *< D*_*ij*,♂_ corresponds to beneficial dominance reversal, in which the lower-valued allele is dominant in females and the higher-valued allele is dominant in males.

### S2.3. Allele frequency dynamics

Our analysis assumes a large population of fixed size *N* under Wright–Fisher population dynamics with mutation and selection (Fisher, 1930; Wright, 1931). We denote the allele frequencies among females and males as *p*_1_, *p*_2_, …, *p*_*n*_ and *q*_1_, *q*_2_, …, *q*_*n*_, respectively, in a population with *n* alleles *x*_1_, …, *x*_*n*_.

Mating is random, and each offspring is assigned a random mother and father from the parent generation. Before viability selection, the expected frequency of an offspring carrying maternally inherited allele *x*_*i*_ and paternally inherited allele *x*_*j*_ is *p*_*i*_*q*_*j*_. Thus, for *i* ≠ *j*, the expected frequency of the unordered heterozygote genotype comprising alleles *x*_*i*_ and *x*_*j*_ is *p*_*i*_*q*_*j*_ + *p*_*j*_*q*_*i*_, while the expected frequency of the homozygote *x*_*i*_*x*_*i*_ is *p*_*i*_*q*_*i*_. Viability selection is sex- and genotype-specific, and the resulting sex ratio can deviate from 1:1. Viability of an individual carrying alleles *x*_*i*_ and *x*_*j*_ is determined by the sex-specific fitness functions *w*_♀_(*x*_*i*_, *a*_*i*,♀_; *x*_*j*_, *a*_*j*,♀_) for females and *w*_♂_(*x*_*i*_, *a*_*i*,♂_; *x*_*j*_, *a*_*j*,♂_) for males (see Eq. S4 and Fig. 1 in the main part). For X-linked loci, the viability of a hemizygous male possessing only a single allele *x*_*i*_ is defined by the function *w*_♂_(*x*_*i*_) (see Eq. S5). Finally, *N* surviving offspring are randomly sampled to represent the adults in the subsequent generation. For our analytical results below, we assume *N* to be sufficiently large for these dynamics to be deterministic, whereas in the simulations (Figs. 3, 4, 5, 6, 7, S3, S4, S5, S6, and S1), stochasticity is introduced by random multinomial sampling of *N* surviving offspring, which constitute the adult population of the next generation.

The expected frequencies of an allele *x*_*i*_ in the subsequent generation after viability selection, collected in the column vector 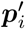 with entries 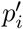 and 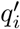, are determined by the matrix recursion 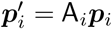, which we write as

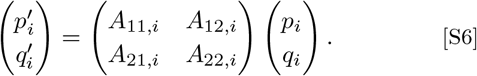

For an autosomal locus, the matrix entries are

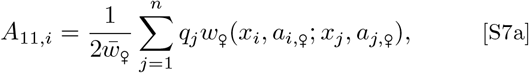

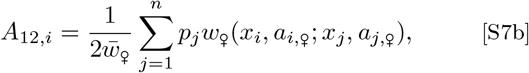

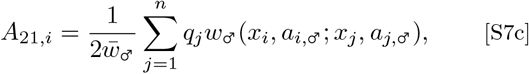

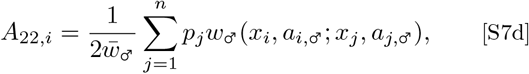

where 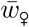 and 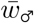 represent the population mean viability among females and males, respectively. Note that *w*_*k*_(*x*_*i*_, *a*_*i,k*_; *x*_*j*_, *a*_*j,k*_) is symmetric in its allelic inputs, i.e., *w*_*k*_(*x*_*i*_, *a*_*i,k*_; *x*_*j*_, *a*_*j,k*_) = *w*_*k*_(*x*_*j*_, *a*_*j,k*_; *x*_*i*_, *a*_*i,k*_).

For an X-linked locus, where males are hemizygous and possess only one allele inherited from their mother, the expected frequency of male offspring carrying allele *x*_*i*_ is directly given by the allele frequency among adult females, *p*_*i*_. Consequently, the entries of the matrix A_*i*_ for an X-linked locus are

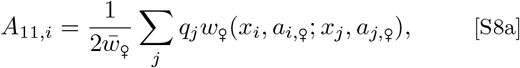

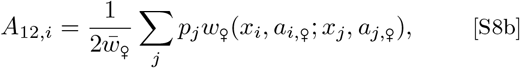

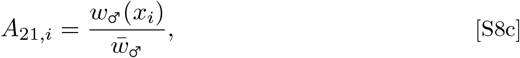

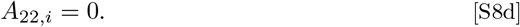

## S3. Evolutionary invasion analysis

We analyse evolution at the focal locus in two steps. The locus affects a quantitative trait through two types of allelic parameters: the allelic value *x* and the sex-specific log-promoter affinities *a*_*i,k*_ (Eq. S1). Importantly, the affinities have no effect in homozygotes, since *z*_*k*_(*x*_*i*_, *a*_*i,k*_; *x*_*i*_, *a*_*i,k*_) = *x*_*i*_. Hence, in a monomorphic population fixed for a single allelic value *x*, any variation in *a* is selectively neutral, and evolution in *a* can only be driven once polymorphism in *x* generates heterozygotes.

Accordingly, we first use adaptive dynamics to analyse gradual evolution in *x* under the assumption that resident affinities are fixed. This yields the singular allelic value *x*^*^ and the conditions under which *x*^*^ is an evolutionary endpoint or an evolutionary branching point where polymorphism in *x* emerges. We then analyse how selection on promoter affinities modifies the subsequent evolutionary dynamics of polymorphic populations.

### S3.1. Selection on promoter affinities

Given polymorphism in *x* (in particular a biallelic polymorphism), selection can act on the log-affinities *a*_*i,k*_. In sex *k*, selection favours changes in affinity that shift the heterozygote phenotype *z*_*k*_ (Eq. S1) toward the sex-specific optimum *θ*_*k*_ (Eq. S4).

This follows by differentiating sex-specific fitness *w*_*k*_ (Eq. S4) with respect to an allele’s sex-specific log-affinity *a*_*i,k*_, which gives

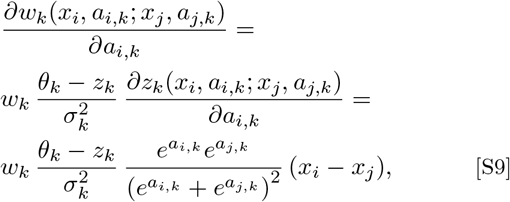

where *w*_*k*_ and *z*_*k*_ without arguments are shorthand for *w*_*k*_(*x*_*i*_, *a*_*i,k*_; *x*_*j*_, *a*_*j,k*_) and *z*_*k*_(*x*_*i*_, *a*_*i,k*_; *x*_*j*_, *a*_*j,k*_), respectively. The second equality uses the derivative of Eq. S1 with respect to *a*_*i,k*_,

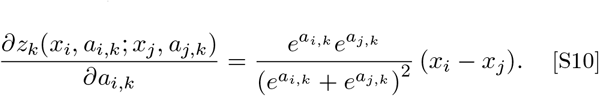

Because *w >* 0, 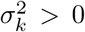, and 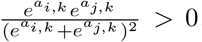, Eq. S9 implies

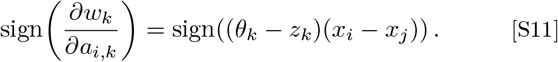

Thus, selection on *a*_*i,k*_ always shifts the heterozygote phenotype toward *θ*_*k*_. Equivalently, in sex *k*, the allele that shifts *z*_*k*_ toward *θ*_*k*_ is selected to increase its affinity (and hence its dominance weight). Consequently, when sex-specific affinities can evolve independently, selection favours beneficial dominance in each sex and thereby generates beneficial sex-specific dominance, which can result in dominance reversal under the independent-affinity model.

### S3.2. Adaptive dynamics of the allelic value *x*

#### S3.2.1. Invasion fitness

In our evolutionary invasion analysis, we consider a large resident population monomorphic for allele *x*, to which we iteratively introduce an initially rare mutant allele *y* = *x* + *ϵ*. In this initial adaptive-dynamics analysis, affinities are held fixed and equal between mutant and resident alleles, and we therefore suppress affinity arguments in the fitness functions below. As long as the mutant allele remains rare, meaning its frequencies *p*_*y*_ and *q*_*y*_ are negligible, the recursion equation for an autosomal locus (Eqs. S6 and S7) simplifies to

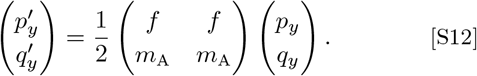

Similarly, for an X-linked locus, the recursion equation (Eqs. S6 and S8) becomes

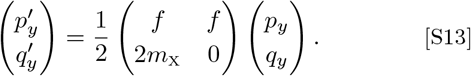

Here, the matrix entry *f* = *w*_♀_(*y, x*)*/w*_♀_(*x, x*) represents the relative fitness of a female carrying the mutant allele, and the entries *m*_A_ = *w*_♂_(*y, x*)*/w*_♂_(*x, x*) and *m*_X_ = *w*_♂_(*y*)*/w*_♂_(*x*) represent the relative fitness of male mutants for autosomal and X-linked loci, respectively.

A rare allele *y* increases in frequency if the dominant eigenvalue *λ*(*y, x*) of the matrix A on the right-hand side of Equations S12 and S13 is larger than one. Following Metz et al. (1992) and Metz (2008), we refer to *λ*(*y, x*) as invasion fitness. This eigenvalue is given by

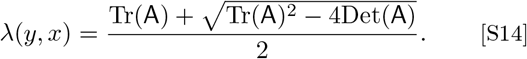

In the autosomal case, we obtain

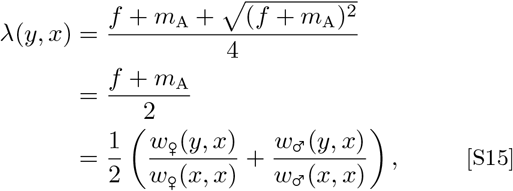

which we recognize as the Shaw-Mohler equation (Shaw and Mohler, 1953).

For an X-linked locus, we obtain

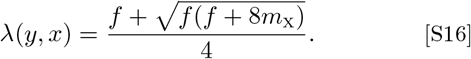

The invasion criterion *λ*(*y, x*) *>* 1 can then be written as

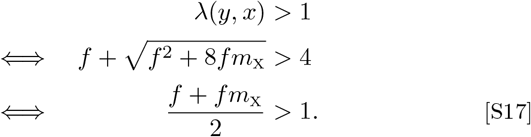

For *f* ≤ 4, the final equivalence follows by isolating the square root and squaring both sides. For *f >* 4, the original invasion criterion is automatically satisfied because *λ*(*y, x*) *> f/*4 *>* 1 and the equivalent condition (*f* + *fm*_X_)*/*2 *>* 1 is likewise automatically satisfied since *m*_X_ *>* 0.

We denote the fitness proxy given by the left-hand side of the last inequality as 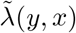. Using the definitions of *f* and *m*_X_, we obtain

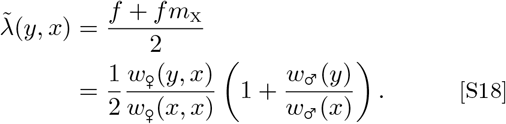

Since 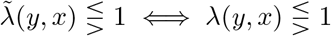, we can use 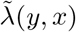 for all further calculations concerning X-linked loci.

#### S3.2.2. Singular points

Given that mutations have small effects (*y* = *x*+*ϵ* with *ϵ* small), the direction of evolutionary change is given by the selection gradient

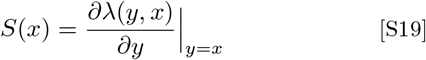

(Geritz et al., 1998). If mutations occur infrequently, a mutant allele *y* either goes extinct or reaches an equilibrium frequency before the next mutant appears. Furthermore, if *S*(*x*) ≠ 0, and still assuming small mutational effects, invasion of *y* implies extinction of *x* (Dercole and Rinaldi, 2008; Priklopil and Lehmann, 2020). Iterating this process results in a trait substitution sequence of allelic values, giving the long-term evolutionary dynamics (Champagnat et al., 2006).

To determine the selection gradient for an autosomal locus, we substitute Equation S15 into Equation S19, resulting in

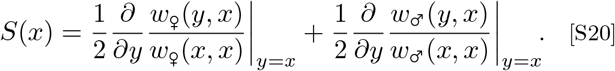

Using the derivatives of the fitness (viability) function provided in Section S6 below, and substituting Equation S51 into Equation S20, we obtain the selection gradient for an autosomal locus,

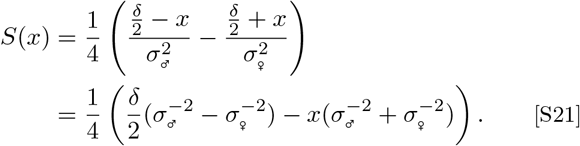

For an X-linked locus, we denote the selection gradient obtained from the invasion fitness in Equation S16 by *S*_X_(*x*). Differentiating Equation S16 with respect to *y* and evaluating at *y* = *x*, where *f* = *m*_X_ = 1, gives

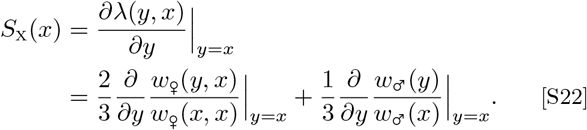

Because the hemizygote derivative is twice as large as the corresponding heterozygote derivative (compare Eqs. S51 and S55), this simplifies to

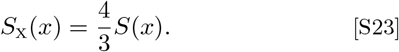

Thus, the autosomal and X-linked selection gradients have the same sign and the same zeros.

The same result can also be obtained from the X-linked invasion-fitness proxy 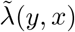. Its gradient is

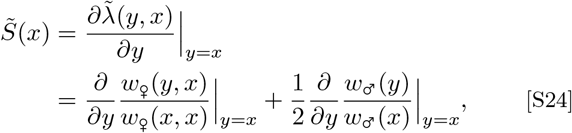

such that

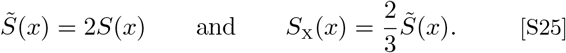

The evolutionary dynamics come to a halt once an allelic value *x*^*^ is reached at which the corresponding selection gradient equals zero. We refer to such allelic values as singular allelic values. We find a unique singular allelic value *x*^*^, which, due to Equation S23, is the same for autosomal and X-linked loci. It is given by

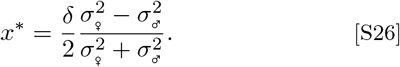

Under symmetric selection strength 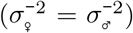, the singular allelic value simplifies to *x*^*^ = 0, representing the midpoint between the two sex-specific optima.

We note that the autosomal selection gradient can be expressed as

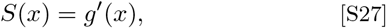

where

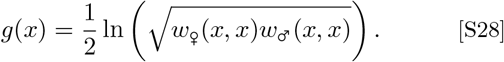

Since *S*_X_(*x*) = 4*S*(*x*)*/*3, evolution acts to increase the same function *g*(*x*) for both autosomal and X-linked loci. Thus, in both cases evolution acts to increase the geometric mean fitness across the two sexes (Charnov, 1982; Leimar, 2001).

#### S3.2.3. Classification of the singular point

Singular allelic values *x*^*^ can be attractors or repellers of trait substitution sequences (Geritz et al., 1998; Doebeli, 2011). Values *x*^*^ that are attractors are referred to as convergence stable. Furthermore, singular allelic values *x*^*^ can be invadable or uninvadable by nearby mutants. Values *x*^*^ that are both convergence stable and uninvadable are evolutionary endpoints (also called continuously stable strategies), whereas values *x*^*^ that are convergence stable and invadable are evolutionary branching points where adaptive allelic polymorphism can emerge.

In our model, the unique singular point *x*^*^ is always convergence stable. This follows directly from

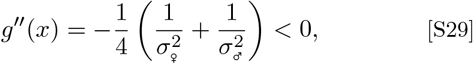

which shows that *g*(*x*) is strictly concave and therefore has its unique maximum at *x*^*^.

The fact that *x*^*^ is always an evolutionary attractor in our model is somewhat sensitive to the functional form of the map from the phenotypic trait *z* to fitness *w* (here given by the Gaussian function in Eq. S4; for a counter-example, see Flintham et al. 2025).

Whether *x*^*^ is an evolutionary endpoint or an evolutionary branching point is therefore determined by its local invadability.

#### S3.2.4. Evolutionary branching under additivity

Whether the evolutionary singular allele *x*^*^ is locally invadable or uninvadable depends on the curvature of invasion fitness at *y* = *x* = *x*^*^. For an autosomal locus, this is determined by the sign of

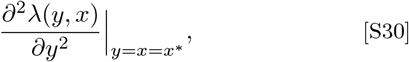

while for an X-linked locus the corresponding criterion is obtained from 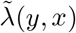. If the relevant second derivative is positive, *x*^*^ is a local fitness minimum and can be invaded by nearby mutants, whereas if it is negative, *x*^*^ is a local fitness maximum and cannot be invaded by nearby mutants. Because *x*^*^ is always convergence stable, a positive second derivative makes it an evolutionary branching point (Geritz et al., 1998; Doebeli, 2011).

We first consider equal promoter affinities, so that alleles act additively, *z*(*x*_*i*_, *x*_*j*_) = (*x*_*i*_ + *x*_*j*_)*/*2. For an autosomal locus, using the second-order derivative given in Equation S58 shows that evolutionary branching occurs if

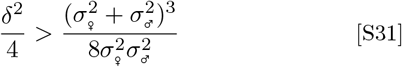

(see Eq. C7 in Flintham et al. 2025 for an identical result). The term on the left-hand side is the variance among the two sex-specific phenotypic optima (recall that *δ* denotes the distance between the optima). Thus, increasing the difference between female and male optima facilitates evolutionary diversification. The right-hand side captures the effect of selection strength and asymmetry. Under symmetric selection, 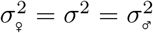, Condition S31 becomes

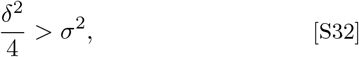

or, equivalently, *δ/σ >* 2. Thus, evolutionary diversification is favoured when selection is sufficiently strong relative to the distance between the sex-specific optima. In this symmetric case, allelic diversification can be interpreted as adaptation to two different environments, namely males and females, and Condition S32 is identical to corresponding branching conditions in models of resource and habitat specialisation (Slatkin, 1979; Dieckmann and Doebeli, 1999; Kisdi and Geritz, 1999; Svardal et al., 2015; Schmid et al., 2024).

For the X-linked case, the corresponding condition is

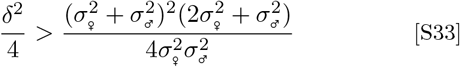

(see Appendix D, in particular Eq. D-20, of Flintham et al. 2025 for the corresponding general X-linked branching condition). Under symmetric selection, this simplifies to *δ*^2^*/σ*^2^ *>* 12, showing that evolutionary branching requires substantially stronger selection for an X-linked locus than for an autosomal locus.

#### S3.2.5. Protected polymorphism near the singular allele

Protected polymorphisms near *x*^*^ can arise under both stabilizing and disruptive selection, but their subsequent evolutionary fate differs (Fig. S7). When *x*^*^ is an evolutionary endpoint, two nearby alleles on opposite sides of *x*^*^ can coexist because heterozygotes have high marginal fitness across the two sexes (marginal overdominance). Selection on the allelic values nevertheless remains stabilizing, so alleles closer to *x*^*^ can subsequently invade and the polymorphism is evolutionarily transient (Fig. S7**A**). Figure 3**i** in the main part shows an individual-based simulation with this outcome, where a protected polymorphism persists for approximately 10^5^ generations after the initial selective sweeps.

When the evolutionary branching condition is fulfilled, by contrast, nearby alleles on opposite sides of *x*^*^ can also coexist, but now experience disruptive selection and diverge away from *x*^*^. In this regime, diversification is associated with marginal homozygote advantage rather than marginal overdominance (see Section S4; (Fig. S7**A**).

We now show that, in the absence of dominance, protected polymorphisms exist in the neighbourhood of *x*^*^ for all parameter values. Specifically, Geritz et al. (1998, Eq. 11) show that in the neighbourhood of *x*^*^ all alleles *x*_1_, *x*_2_ with *x*_1_ *< x*^*^ *< x*_2_ that are equidistant from *x*^*^ (|*x*_1_ − *x*^*^| = |*x*_2_ − *x*^*^|) can coexist if and only if

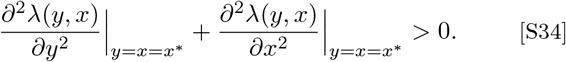

For an autosomal locus, using the second-order derivatives of the invasion fitness function (Eqs. S58 and S60), we obtain

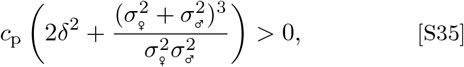

where *c*_p_ is a positive constant defined in Equation S59. Hence, this condition always holds.

For an X-linked locus, we use the fitness proxy 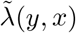. Since we consider the absence of dominance here, we set *d*_♀_ = 0 in the second-order mutant and resident derivatives (Eqs. S72 and S74). Their sum is

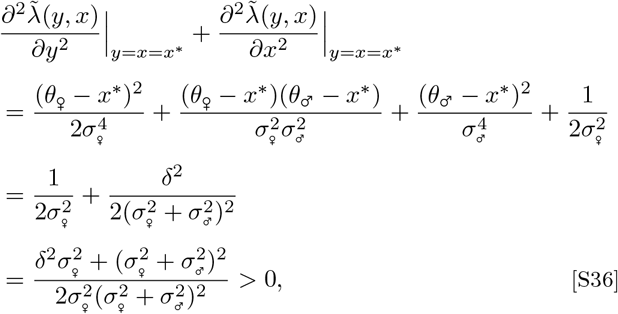

where the second equality follows from Equation S26, which gives 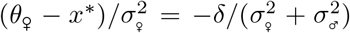 and 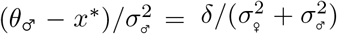. Thus, the condition for protected polymorphism near *x*^*^ is always fulfilled for X-linked loci as well.

This local coexistence can be understood in relation to classical two-allele models of sexually antagonistic selection (Kidwell et al., 1977). Gradual evolution leads the population toward *x*^*^, where the opposing female and male contributions to the selection gradient balance. Thus, even when selection differs in strength between the sexes, evolution toward *x*^*^ brings the population into a region where small allelic deviations on either side of *x*^*^ generate opposing, balanced sex-specific fitness effects. When *x*^*^ is an evolutionary endpoint, this balance generates marginal overdominance between nearby alleles on opposite sides of *x*^*^, allowing them to mutually invade even though selection on their allelic values remains locally convergent. When the branching condition is fulfilled, the same local mutual invasibility instead occurs under disruptive selection, causing the coexisting allelic lineages to subsequently diverge.

#### S3.2.6. Coevolution between the trait locus and its sex-specific affinities

We now generalize the additive branching conditions by allowing the two coexisting allelic lineages to differ in their sex-specific promoter affinities. Below, we show that the effect of beneficial sex-specific dominance on disruptive selection near *x*^*^ scales inversely with the allelic difference in *x*. Consequently, when a BAP emerges between alleles that have similar *x*-values, only a very small difference in promoter affinities is required to turn selection disruptive. Once disruptive selection maintains and separates the two allelic lineages, this can initiate a coevolutionary feedback: the affinities (*a*_♀_, *a*_♂_) evolve toward increased beneficial sex-specific domina facilitating further divergence in *x*, which in turn strengthens selection on promoter affinities. This change from convergent to divergent evolution of the two allelic lineages is illustrated in Figure S7, where beneficial sex-specific dominance reverses the local direction of selection acting on a protected polymorphism around *x*^*^: under parameter combinations where selection is stabilizing in the additive case, selection can turn disruptive once beneficial sex-specific dominance is sufficiently strong.

As above, when comparing two alleles we index them such that *x*_*i*_ *> x*_*j*_. We define

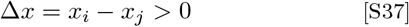

and their sex-specific difference in log-affinity as

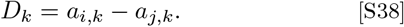

For the mathematical analysis below, we express this affinity difference relative to the divergence in allelic values as

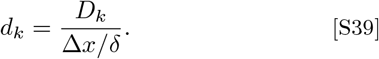

Thus, *d*_*k*_ measures the difference in log-affinity between the two alleles relative to their divergence in allelic value Δ*x*, scaled by the distance *δ* between the sex-specific optima. Because Δ*x >* 0, *d*_*k*_ and *D*_*k*_ have the same sign: positive values imply dominance of the allele with the larger allelic value, whereas negative values imply dominance of the allele with the smaller allelic value.

For any given pair of alleles, their log-affinities can equivalently be represented by a linear function of allelic values,

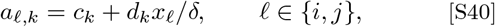

where *c*_*k*_ is a constant. This representation introduces no additional assumption for the two focal alleles: choosing *d*_*k*_ according to Equation S39, together with the corresponding *c*_*k*_ exactly reproduces their observed difference in log-affinity. The parameter *d*_*k*_ can therefore be interpreted as the local change in log-affinity associated with a change in allelic value, which is useful when evaluating selection on nearby allelic lineages.

Substituting this local representation into Equation S1 and cancelling the common factor 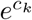 gives the composite phenotype function

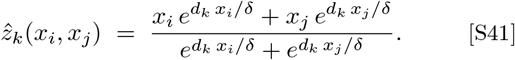

We use a hat to distinguish this composite function from the original phenotype function in Equation S1. We denote by *ŵ*_*k*_ the viability function obtained by substituting 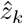 into Equation S4, and by 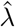 the corresponding autosomal invasion fitness obtained from Equation S15. When *D*_♀_ = *D*_♂_ = 0, the two focal alleles have equal promoter affinities within each sex, and 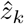 reduces to the additive phenotype.

##### Autosomal locus

For an autosomal locus, we obtain

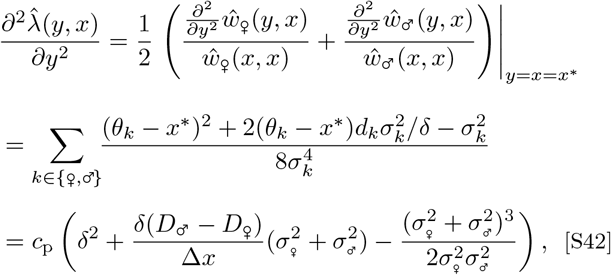

where *c*_p_ is a positive constant defined in Equation S59. The second equality is obtained using the second-order partial derivative of *ŵ*_*k*_ with respect to *y*, as given in Equation S68, and the last step follows from using the expression for *x*^*^ given by Equation S26 and *d*_*k*_ = *δD*_*k*_*/*Δ*x*.

In summary, the condition for disruptive selection at *x*^*^ is

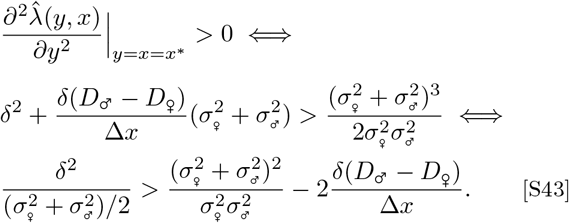

Thus, beneficial sex-specific dominance (*D*_♂_ *> D*_♀_) facilitates allelic divergence, whereas detrimental sex-specific dominance (*D*_♂_ *< D*_♀_) inhibits it. In the absence of sex-specific dominance (*D*_♂_ = *D*_♀_), Condition S43 reduces to the condition for evolutionary branching under additivity or parallel dominance (Condition S31), matching Eq. C7 of Flintham et al. (2025). Assuming symmetry (*σ*^2^ = *σ*^2^ = *σ*^2^), Condition S43 can be rewritten as

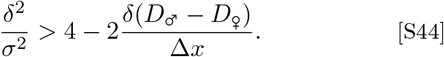

Under symmetry and in the absence of sex-specific dominance, Condition S44 simplifies further to *δ/σ >* 2.

##### X-linked locus

For an X-linked locus, an analogous calculation results in

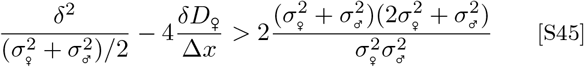

as the condition for disruptive selection at *x*^*^. The terms in this condition are analogous to the corresponding terms in Condition S43. Because only females are diploid at an X-linked locus, dominance affects the condition for disruptive selection only through females. Female-beneficial dominance (*D*_♀_ *<* 0) facilitates evolutionary diversification, whereas *D*_♀_ *>* 0 inhibits it. Under symmetry 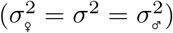, Condition S45 simplifies to

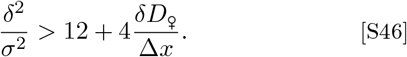

In the absence of dominance (*D*_♀_ = 0), this condition further simplifies to *δ*^2^*/σ*^2^ *>* 12, recovering Condition S33.

### S3.3. Large mutational steps

When the evolutionary branching condition (Condition S31 for autosomal loci or Condition S33 for X-linked loci) is not fulfilled, the singular allele *x*^*^ is a local fitness maximum and therefore uninvadable by mutants of small effect. Under the standard small-step assumption of adaptive dynamics, *x*^*^ is then an evolutionary endpoint. However, *x*^*^ need not be a global fitness maximum and may still be invadable by mutants with larger phenotypic effects.

To assess global invadability, we numerically computed

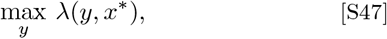

for autosomal loci, and the corresponding quantity 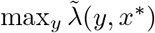 for X-linked loci. We then tested whether the respective maximum exceeds one. If so, a resident population at *x*^*^ can be invaded by a mutant of sufficiently large effect. Otherwise, *x*^*^ is globally uninvadable.

The resulting thresholds are shown by the dotted white lines in Figure 6 and Figure 7**A**. Allowing larger mutational steps can therefore expand the parameter region in which polymorphism arises via invasion of *x*^*^, even when small-effect mutants cannot invade. This effect is especially prominent for X-linked loci (Fig. 6**B**). Figure S8 contrasts a globally uninvadable autosomal singular allele with an X-linked singular allele that is locally uninvadable by nearby mutants but invadable by mutants of sufficiently large effect.

## S4. Geometrical interpretation

The condition for evolutionary branching at an autosomal locus in the absence of sex-specific dominance (*D*_♂_ = *D*_♀_) has a useful geometric interpretation. For this, we note that the growth rate of a rare mutant allele *y* invading a resident population in which all individuals are homozygous for an allele *x* is given by the Shaw-Mohler equation (Eq. S15), which is a linear combination of the female and male fitnesses of individuals heterozygous for *y* and *x*. A consequence of this linearity (Levins, 1968; Charnov, 1982; Rueffler et al., 2004) is that a population monomorphic for the intermediate allelic value *x*^*^ is uninvadable if the red curve shown in Figure S9 is concave at *z*^*^ (negative second derivative, as shown in the first two panels), resulting in stabilizing selection. Conversely, a population monomorphic for *x*^*^ is invadable if this curve is convex at *z*^*^ (positive second derivative, third panel), resulting in evolutionary branching due to disruptive selection. A further consequence is that, in a population with two segregating alleles *x*_1_ *< x*^*^ *< x*_2_ that emerged from evolutionary branching, the fitness of homozygotes—when averaged over the two sexes—is higher than the average fitness of heterozygotes (marginal underdominance). In contrast, if the allele *x*^*^ experiences stabilizing selection, this is because the fitness of individuals homozygous for this allele, when averaged over females and males, is higher than that of alleles more specialized for one sex.

The solid white curves in Figure 6**A** show the combinations of female and male selection strength at which the red curves in Figure S9 change from locally concave to convex at *z*^*^. Consequently, for an autosomal locus in the absence of sex-specific dominance, allelic diversification is possible only above and to the right of the solid white curves, in the limit of small mutational effects. The alleles in the resulting BAP are maintained by balancing selection even though the fitness of homozygous individuals, when averaged over the two sexes, is greater than that of heterozygotes. The corresponding X-linked branching condition is more stringent because X-linked inheritance weights female and male selection differently (Condition S33); it therefore cannot be inferred from the curvature of the same Shaw-Mohler fitness trade-off alone.

Condition S43 shows that, for autosomal loci, the parameter region allowing allelic diversification expands with increasing beneficial sex-specific dominance (*D*_♂_ − *D*_♀_ *>* 0). This is shown in Figure 7**B**–**E**, where polymorphism can emerge across the full parameter space. An analogous expansion occurs for X-linked loci through female-beneficial dominance (Condition S45; Fig. 7**F**). To understand this, we again make use of a geometric argument. Under sex-specific dominance, heterozygous females and males carrying the same two alleles express different trait values *z*. Thus, the combination of female and male fitnesses of such heterozygotes no longer lies on the red curves in Figure S9, but instead lies above them if sex-specific dominance is beneficial and below them if it is detrimental. As an example, the filled grey dots indicate female and male fitness under full beneficial dominance reversal (*D*_♀_ large and negative and *D*_♂_ large and positive) for individuals heterozygous for alleles coding for the female and male optimal phenotypes. Thus, in a population with two segregating alleles *x*_1_ *< x*^*^ *< x*_2_ that code for the two sex-specific optimal phenotypes, or phenotypes between these optima, the fitness of homozygotes, when averaged over the two sexes, is lower than the average fitness of heterozygotes (marginal overdominance): heterozygous individuals enjoy an average fitness that no single allele can achieve in homozygous individuals. Adaptive allelic diversification in the presence of sex-specific dominance for combinations of female and male selection strengths that lie below and to the left of the solid white curves in Figure 7**B**–**E** is therefore driven by marginal heterozygote advantage.

## S5. The distribution of allelic values

Intuitively, one might expect evolution to result in allelic values distributed between the sex-specific optima. However, under additivity, the evolutionary endpoint consists of two alleles with asymmetric allelic values (see Fig. 4**iii**). One allele, occurring at frequency 3*/*4, codes for the optimal phenotype in one sex, while the other allele, occurring at frequency 1*/*4, overshoots the optimum of the other sex. For the parameterization used in our figures (*δ* = 1), this results in two allelic values differing by 2 rather than 1, the distance between the sex-specific optima (see lower panels in Fig. 6). This asymmetry generates a frequency of 1*/*2 for high-marginal-fitness genotypes (assuming strong selection), rather than the frequency of 1*/*4 that would be achieved by two alleles each coding for one sex-specific optimum. An analogous result was found by Kisdi and Geritz (1999) in a model studying habitat specialization.

Under the independent-affinity model, coexisting alleles occur between the sex-specific optima (Figs. 3**ii, iib, iic**, 4**iv**, S1**v**, S2). By contrast, under the shared-promoter model, the distribution of allelic values can be highly skewed and extend beyond the sex-specific optima (Figs. S1**vii,viii** and S2). For X-linked loci, when selection is stronger in males than in females, several coexisting alleles can likewise occupy only a narrow range of trait space (orange and red regions in Fig. 7**F** but blue regions in the corresponding panel of Fig. S2), similar to the sex-limited dominance scenario in Fig. 7**C**.

## S6. Derivatives

In this section, we present derivatives of the phenotypic trait function *z*_*k*_ (Eq. S1), the viability functions *w*_*k*_ (Eqs. S4 and S5), and the invasion fitness functions (Eqs. S15 and S18) used in the calculations detailed above.

### S6.1. Derivatives of the phenotype function

The partial derivative of the phenotype function *z*_*k*_(*y, a*_*y,k*_; *x, a*_*x,k*_) with respect to one allelic value *y* equals

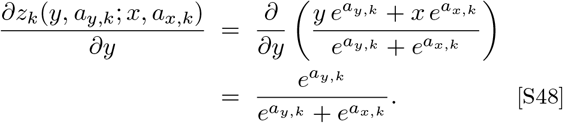

When evaluated at *a*_*y,k*_ = *a*_*x,k*_ = *a*_*k*_, Eq. S48 simplifies to

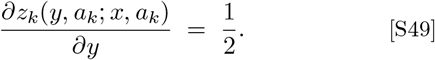

The second partial derivative is

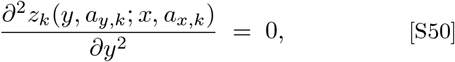

because the first-order derivative (Eq. S48) is independent of *y* when the affinities are held fixed.

### S6.2. Derivatives of the fitness (viability) functions

#### S6.2.1. First partial mutant derivative

The partial derivative of the fitness function with respect to *y* equals

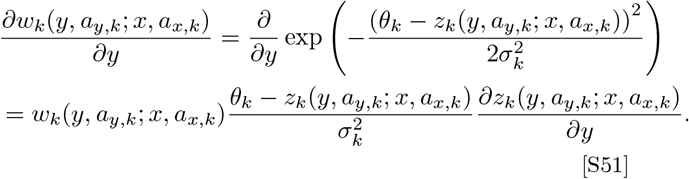

When evaluated at (*y, a*_*y,k*_) = (*x, a*_*x,k*_), this becomes

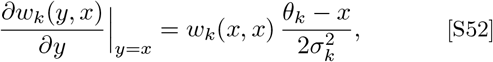

where we use that *z*_*k*_(*x, x*) = *x* and Equations S4 and S49.

#### S6.2.2. Second partial mutant derivative

The second partial derivative of *w*_*k*_ with respect to *y*, evaluated at (*y, a*_*y,k*_) = (*x, a*_*x,k*_) with equal affinities, can be derived from Equation S51:

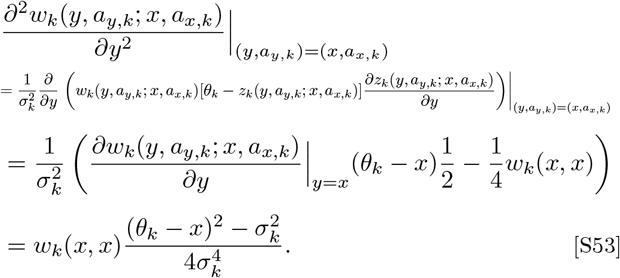

Here we used Eqs. S49, S50, S51, and S52.

#### S6.2.3. Homozygote/hemizygote derivative

For both homozygotes and hemizygotes, the phenotype and thus fitness function is independent of the affinities. The first derivative of the fitness function of a homozygote *w*_*k*_(*x, x*) with respect to *x* is

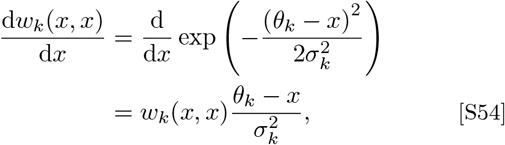

and similarly for a hemizygote male,

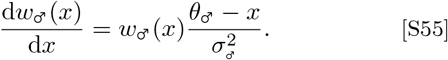

#### S6.2.4. Second homozygote/hemizygote derivative

The second derivative of *w*_*k*_(*x, x*) equals

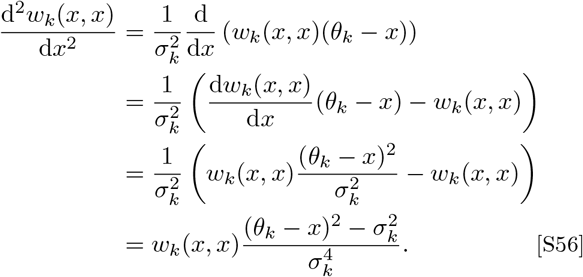

Similarly, for a hemizygote male,

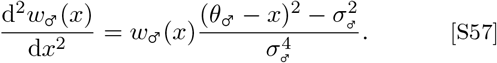

### S6.3. Derivatives of autosomal invasion fitness

#### S6.3.1. Second partial mutant derivative

The second partial derivative of the autosomal invasion fitness *λ*(*y, x*) with respect to the mutant allelic value *y*, evaluated at the singular point, is

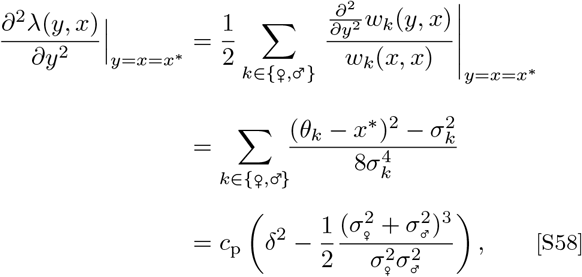

where

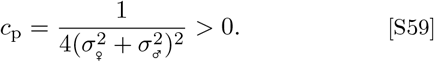

#### S6.3.2. Second resident derivative

For the second derivative with respect to the resident allelic value *x*,

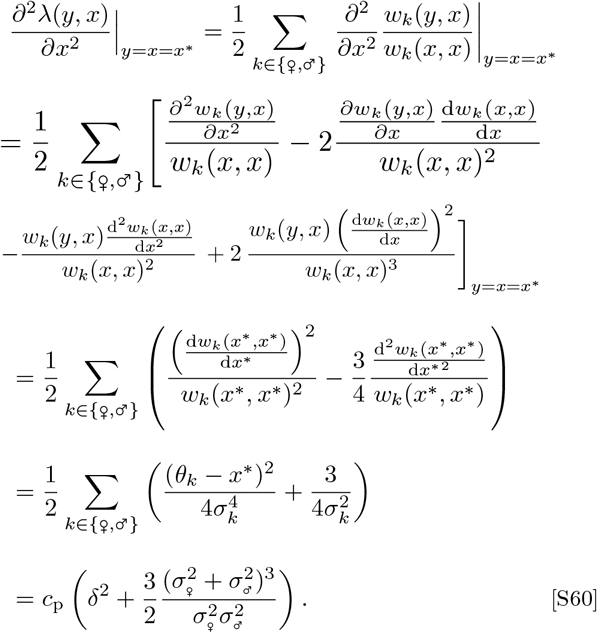

Here we used

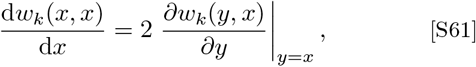

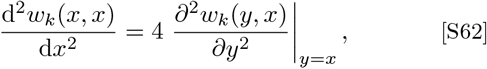

and Eqs. S51, S53, S54, and S56.

## S7. The reparameterised phenotype function

### S7.1. Derivatives of the phenotype function

For the local analysis, the affinity difference between the two alleles is given by *D*_*k*_(*y, x*) = *d*_*k*_(*y* − *x*)*/δ*. Factoring out the common term 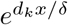, we can write

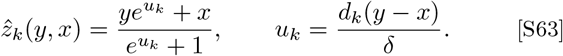

Since *∂u*_*k*_*/∂y* = *d*_*k*_*/δ*, the quotient rule gives

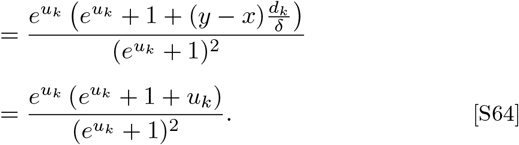

When evaluated at *y* = *x* (and hence *u*_*k*_ = 0), Eq. S64 simplifies to

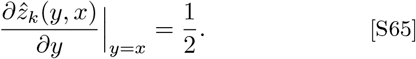

The second partial derivative, evaluated at *y* = *x*, is

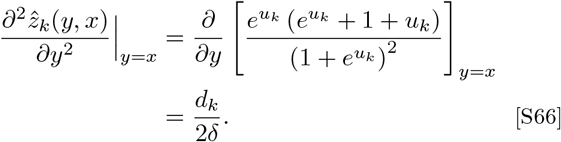

### S7.2. Derivatives of the reparameterised fitness (viability) functions

#### S7.2.1. First partial mutant derivative

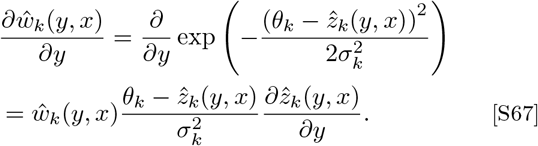

#### S7.2.2. Second partial mutant derivative

The second partial derivative of 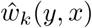 with respect to *y*, evaluated at *y* = *x*, can be derived from Eq. S67:

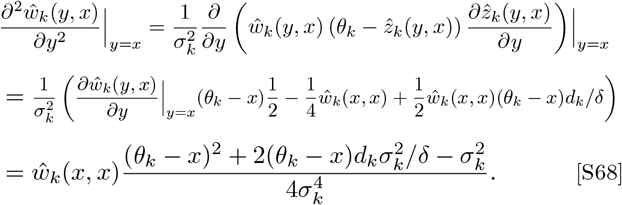

Here we used Eqs. S67, S65, and S66.

### S7.3. Derivatives of the X-linked invasion-fitness proxy

For an X-linked locus, we write the invasion-fitness proxy (Eq. S18) as

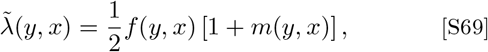

where

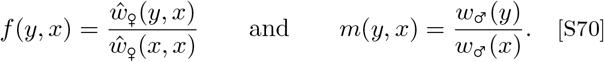

At *y* = *x*, the derivatives with respect to the mutant allelic value *y* are

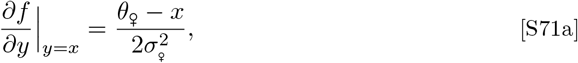

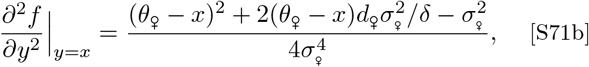

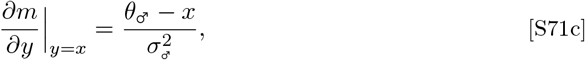

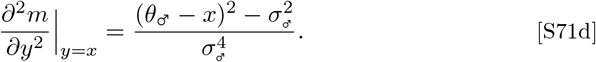

These expressions follow from Eqs. S67, S65, S68, S55, and S57.

Differentiating 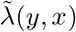 twice with respect to *y* gives

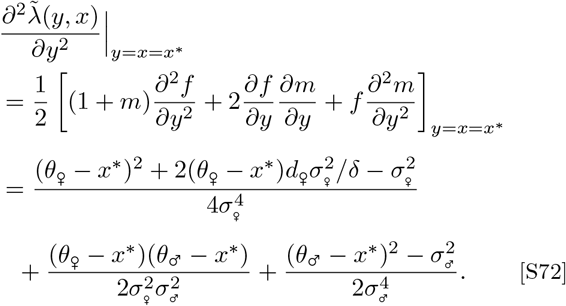

At *y* = *x*, the corresponding derivatives with respect to the resident allelic value *x* are

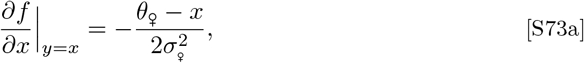

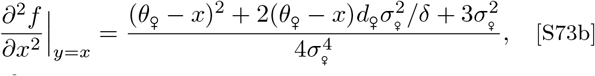

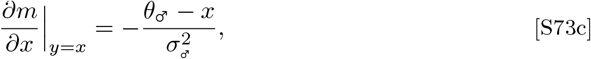

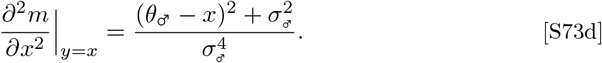

For the female terms, symmetry of *ŵ*_♀_(*y, x*) implies that the partial derivatives of its numerator with respect to *x* at *y* = *x* equal the corresponding partial derivatives with respect to *y*. Combining these with quotient differentiation and the homozygote derivatives in Eqs. S54 and S56 gives the first two expressions in Eqs. S73. The male terms follow directly from Eqs. S55 and S57.

Differentiating 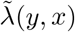 twice with respect to *x* gives

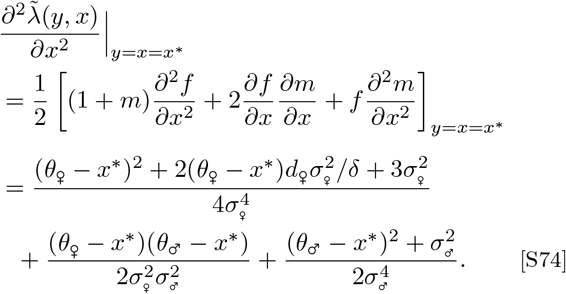

## References

Arnqvist, G. 2011. Assortative mating by fitness and sexually antagonistic genetic variation. Evolution, 65:2111–2116.

Arnqvist, G., and L. Rowe. 2005. Sexual conflict, vol. 31. Princeton university press.

Arnqvist, G., N. Vellnow, and L. Rowe. 2014. The effect of epistasis on sexually antagonistic genetic variation. Proceedings of the Royal Society B: Biological Sciences, 281:20140489.

Barson, N. J., T. Aykanat, K. Hindar, M. Baranski, G. H. Bolstad,P. Fiske, C. Jacq, A. J. Jensen, S. E. Johnston, S. Karlsson, et al. 2015. Sex-dependent dominance at a single locus maintains variation in age at maturity in salmon. Nature, 528:405.

Bergeron, L. A., S. Besenbacher, J. Zheng, P. Li, M. F. Bertelsen,B. Quintard, J. I. Hoffman, Z. Li, J. St. Leger, C. Shao, et al. 2023. Evolution of the germline mutation rate across vertebrates. Nature, 615:285–291.

Billiard, S., and V. Castric. 2011. Evidence for fisher’s dominance theory: how many ‘special cases’? Trends in Genetics, 27:441–445.

Billiard, S., V. Castric, and V. Llaurens. 2021. The integrative biology of genetic dominance. Biological Reviews, 96:2925–2942.

Bonduriansky, R. 2007. The genetic architecture of sexual dimorphism: The potential roles of genomic imprinting and condition-dependence. Pages 176–184 in D. J. Fairbairn, W. U. Blanckenhorn, and T. Székely, eds. Sex, Size and Gender Roles: Evolutionary Studies of Sexual Size Dimorphism. Oxford University Press.

Bonduriansky, R., and S. F. Chenoweth. 2009. Intralocus sexual conflict. Trends in ecology & evolution, 24:280–288.

Brud, E. 2025. Season-specific dominance broadly stabilizes polymorphism under symmetric and asymmetric multivoltinism. Genetics, 229:iyaf028.

Charlesworth, B. 2015. Causes of natural variation in fitness: evidence from studies of drosophila populations. Proceedings of the National Academy of Sciences, 112:1662–1669.

Charlesworth, B., and K. A. Hughes. 2000. The maintenance of genetic variation in life-history traits. Pages 369–392 in Evolutionary genetics: from molecules to morphology. Cambridge University Press.

Charnov, E. L. 1982. The Theory of Sex Allocation, vol. 18. Princeton University Press.

Chen, J., V. Nolte, and C. Schlötterer. 2015. Temperature stress mediates decanalization and dominance of gene expression in drosophila melanogaster. PLoS Genetics, 11:e1004883.

Connallon, T., and S. F. Chenoweth. 2019. Dominance reversals and the maintenance of genetic variation for fitness. PLoS Biology, 17:e3000118.

Connallon, T., and A. G. Clark. 2012. A general population genetic framework for antagonistic selection that accounts for demography and recurrent mutation. Genetics, 190:1477–1489.

Connallon, T., and A. G. Clark. 2014. Balancing selection in species with separate sexes: insights from fisher’s geometric model. Genetics, 197:991–1006.

Connallon, T., and L. L. Knowles. 2005. Intergenomic conflict revealed by patterns of sex-biased gene expression. Trends in Genetics, 21:495–499.

Connallon, T., and G. Matthews. 2019. Cross-sex genetic correlations for fitness and fitness components: connecting theoretical predictions to empirical patterns. Evolution Letters, 3:254–262.

Curtsinger, J. W., P. M. Service, and T. Prout. 1994. Antagonistic pleiotropy, reversal of dominance, and genetic polymorphism. The American Naturalist, 144:210–228.

Dean, R., and J. E. Mank. 2014. The role of sex chromosomes in sexual dimorphism: discordance between molecular and phenotypic data. Journal of Evolutionary Biology, 27:1443–1453.

Doebeli, M. 2011. Adaptive Diversification (MPB-48). Princeton University Press.

Ellegren, H., and J. Parsch. 2007. The evolution of sex-biased genes and sex-biased gene expression. Nature Reviews Genetics, 8:689–698.

Fairbairn, D. J., D. A. Roff, and M. E. Wolak. 2023. Tests for associations between sexual dimorphism and patterns of quantitative genetic variation in the water strider, aquarius remigis. Heredity, 131:109–118.

Fan, H., and J.-Y. Chu. 2007. A brief review of short tandem repeat mutation. Genomics, proteomics & bioinformatics, 5:7–14.

Fisher, R. A. 1930. The genetical theory of natural selection. Oxford University Press.

Fisher, R. A. 1931. The evolution of dominance. Biological Reviews of the Cambridge Philosophical Society, 6:345–368.

Flintham, E. 2025. The evolution of sex-specific gene expression in polygenic traits. Journal of evolutionary biology, page voaf050.

Flintham, E., T. Lesaffre, S. Otto, and C. Mullon. 2026. Sexually antagonistic selection: a review of the theory and its implications. EcoEvoRxiv.

Flintham, E., V. Savolainen, S. P. Otto, M. Reuter, and C. Mullon. 2025. The maintenance of genetic polymorphism underlying sexually antagonistic traits. Evolution Letters, page qrae059.

Fry, J. D. 2010. The genomic location of sexually antagonistic variation: some cautionary projects. Evolution, 64:1510–1516.

Gavrilets, S., and D. Waxman. 2002. Sympatric speciation by sexual conflict. Proceedings of the National Academy of Sciences, 99:10533–10538.

Geeta Arun, M., A. Agarwala, Z. A. Syed, M. Kashyap, S. Venkatesan,T. S. Chechi, V. Gupta, and N. G. Prasad. 2021. Experimental evolution reveals sex-specific dominance for surviving bacterial infection in laboratory populations of drosophila melanogaster. Evolution Letters, 5:657–671.

Geritz, S. A. H., É. Kisdi, G. Meszéna, and J. A. J. Metz. 1998. Evolutionarily singular strategies and the adaptive growth and branching of the evolutionary tree. Evolutionary Ecology, 12:35–57.

Gibson, J. R., A. K. Chippindale, and W. R. Rice. 2002. The x chromosome is a hot spot for sexually antagonistic fitness variation. Proceedings of the Royal Society of London. Series B: Biological Sciences, 269:499–505.

Glaser-Schmitt, A., and J. Parsch. 2018. Functional characterization of adaptive variation within a cis-regulatory element influencing drosophila melanogaster growth. PLoS Biology, 16:e2004538.

Glaser-Schmitt, A., M. J. Wittmann, T. J. Ramnarine, and J. Parsch. 2021. Sexual antagonism, temporally fluctuating selection, and variable dominance affect a regulatory polymorphism in Drosophila melanogaster. Molecular Biology and Evolution, 38:4891–4907.

Grieshop, K., and G. Arnqvist. 2018. Sex-specific dominance reversal of genetic variation for fitness. PLoS biology, 16:e2006810.

Grieshop, K., E. K. Ho, and K. R. Kasimatis. 2024. Dominance reversals: The resolution of genetic conflict and maintenance of genetic variation. Proceedings of the Royal Society B: Biological Sciences, 291:20232816.

Haldane, J. B. S. 1956. The theory of selection for melanism in lepidoptera. Proceedings of the Royal Society of London. Series B, Biological Sciences, 145:303–306.

Hawkes, M., C. Gamble, E. Turner, M. Carey, N. Wedell, andD. Hosken. 2016. Intralocus sexual conflict and insecticide resistance. Proceedings of the Royal Society B: Biological Sciences, 283:20161429.

Haygood, R. 2004. Sexual conflict and protein polymorphism. Evolution, 58:1414–1423.

Hedrick, P. W. 1999. Antagonistic pleiotropy and genetic polymorphism: a perspective. Heredity, 82:126–133.

Hollis, B., D. Houle, Z. Yan, T. J. Kawecki, and L. Keller. 2014. Evolution under monogamy feminizes gene expression in drosophila melanogaster. Nature communications, 5:3482.

Huber, C. D., A. Durvasula, A. M. Hancock, and K. E. Lohmueller. 2018. Gene expression drives the evolution of dominance. Nature communications, 9:2750.

Innocenti, P., and E. H. Morrow. 2010. The sexually antagonistic genes of drosophila melanogaster. PLoS biology, 8:e1000335.

Kaiser, V. B., L. Talmane, Y. Kumar, F. Semple, M. MacLennan,D. D. D. Study, D. R. FitzPatrick, M. S. Taylor, and C. A. Semple. 2021. Mutational bias in spermatogonia impacts the anatomy of regulatory sites in the human genome. Genome Research, 31:1994–2007.

Karageorgi, M., A. S. Lyulina, M. C. Bitter, E. Lappo, S. I. Greenblum,Z. K. Mouza, C. T. Tran, A. V. Huynh, H. Oken, P. Schmidt, andD. A. Petrov. 2024. Dominance reversal protects large-effect resistance polymorphisms in temporally varying environments. bioRxiv, pages 2024–10.

Kaufmann, P., J. L. Rönn, E. Immonen, and G. Arnqvist. 2024. Sex-specific dominance of gene expression in seed beetles. Molecular biology and evolution, 41:msae244.

Khudiakova, K. A., N. H. Barton, and G. Arnqvist. 2025. Sign epistasis extends the effects of balancing selection on genetic diversity. bioRxiv, pages 2025–04.

Kidwell, J., M. Clegg, F. Stewart, and T. Prout. 1977. Regions of stable equilibria for models of differential selection in the two sexes under random mating. Genetics, 85:171–183.

Kimura, M., and J. F. Crow. 1964. The number of alleles that can be maintained in a finite population. Genetics, 49:725–738.

Kisdi, É., and S. A. H. Geritz. 1999. Adaptive dynamics in allele space: evolution of genetic polymorphism by small mutations in a heterogeneous environment. Evolution, pages 993–1008.

Kopp, M., and J. Hermisson. 2006. The evolution of genetic architecture under frequency-dependent disruptive selection. Evolution, 60:1537–1550.

Leimar, O. 2001. Evolutionary change and darwinian demons. Selection, 2:65–72.

Lewontin, R., L. Ginzburg, and S. Tuljapurkar. 1978. Heterosis as an explanation for large amounts of genic polymorphism. Genetics, 88:149–169.

Liao, X., W. Zhu, J. Zhou, H. Li, X. Xu, B. Zhang, and X. Gao. 2023. Repetitive dna sequence detection and its role in the human genome. Communications biology, 6:954.

Lonn, E., E. Koskela, T. Mappes, M. Mokkonen, A. M. Sims, andP. C. Watts. 2017. Balancing selection maintains polymorphisms at neurogenetic loci in field experiments. Proceedings of the National Academy of Sciences, 114:3690–3695.

Lynch, M., F. Ali, T. Lin, Y. Wang, J. Ni, and H. Long. 2023. The divergence of mutation rates and spectra across the tree of life. The EMBO Reports, 24:EMBR202357561.

Manna, F., G. Martin, and T. Lenormand. 2011. Fitness landscapes: an alternative theory for the dominance of mutation. Genetics, 189:923–937.

Meiklejohn, C. D., J. D. Coolon, D. L. Hartl, and P. J. Wittkopp. 2014. The roles of cis-and trans-regulation in the evolution of regulatory incompatibilities and sexually dimorphic gene expression. Genome Research, 24:84–95.

Mérot, C., V. Llaurens, E. Normandeau, L. Bernatchez, and M. Wellenreuther. 2020. Balancing selection via life-history trade-offs maintains an inversion polymorphism in a seaweed fly. Nature communications, 11:670.

Metz, J. A. J., R. M. Nisbet, and S. A. H. Geritz. 1992. How should we define ‘fitness’ for general ecological scenarios? Trends in Ecology & Evolution, 7:198–202.

Mishra, P., T. S. Barrera, K. Grieshop, and A. F. Agrawal. 2022. Cis-regulatory variation in relation to sex and sexual dimorphism in drosophila melanogaster. bioRxiv, 2022.09.20.508724.

Mitchell-Olds, T., J. H. Willis, and D. B. Goldstein. 2007. Which evolutionary processes influence natural genetic variation for phenotypic traits? Nature Reviews Genetics, 8:845–856.

Niitepõld, K., and M. Saastamoinen. 2017. A candidate gene in an ecological model species: Phosphoglucose isomerase (pgi) in the glanville fritillary butterfly (melitaea cinxia). Pages 259–273 in Annales Zoologici Fennici. Vol. 54(1–4). BioOne.

Otto, S. P., and D. Bourguet. 1999. Balanced polymorphisms and the evolution of dominance. The American Naturalist, 153:561–574.

Pamilo, P. 1979. Genic variation at sex-linked loci: quantification of regular selection models. Hereditas, 91:129–133.

Patten, M. M., and D. Haig. 2009. Maintenance or loss of genetic variation under sexual and parental antagonism at a sex-linked locus. Evolution, 63:2888–2895.

Pearse, D. E., N. J. Barson, T. Nome, G. Gao, M. A. Campbell,A. Abadía-Cardoso, E. C. Anderson, D. E. Rundio, T. H. Williams,K. A. Naish, et al. 2019. Sex-dependent dominance maintains migration supergene in rainbow trout. Nature ecology & evolution, 3:1731–1742.

Posavi, M., G. W. Gelembiuk, B. Larget, and C. E. Lee. 2014. Testing for beneficial reversal of dominance during salinity shifts in the invasive copepod eurytemora affinis, and implications for the maintenance of genetic variation. Evolution, 68:3166–3183.

Prout, T. 2000. How well does opposing selection maintain variation. Pages 157–181 in Evolutionary genetics: from molecules to morphology. Cambridge University Press.

Puixeu, G., A. Macon, and B. Vicoso. 2023. Sex-specific estimation of cis and trans regulation of gene expression in heads and gonads of drosophila melanogaster. G3: Genes, Genomes, Genetics, 13:jkad121.

Reinhold, K. 1998. Sex linkage among genes controlling sexually selected traits. Behavioral Ecology and Sociobiology, 44:1–7.

Rice, W. R. 1984. Sex chromosomes and the evolution of sexual dimorphism. Evolution, 38:735–742.

Rice, W. R., and A. K. Chippindale. 2002. The evolution of hybrid infertility: perpetual coevolution between gender-specific and sexually antagonistic genes. Genetics of mate choice: from sexual selection to sexual isolation, pages 179–188.

Rogers, T. F., D. H. Palmer, and A. E. Wright. 2021. Sex-specific selection drives the evolution of alternative splicing in birds. Molecular Biology and Evolution, 38:519–530.

Rowe, L., S. F. Chenoweth, and A. F. Agrawal. 2018. The genomics of sexual conflict. The American Naturalist, 192:274–286.

Rusuwa, B. B., H. Chung, S. L. Allen, F. D. Frentiu, and S. F. Chenoweth. 2022. Natural variation at a single gene generates sexual antagonism across fitness components in drosophila. Current Biology, 32:3161–3169.

Ruzicka, F., and T. Connallon. 2020. Is the x chromosome a hot spot for sexually antagonistic polymorphisms? biases in current empirical tests of classical theory. Proceedings of the Royal Society B: Biological Sciences, 287:20201869.

Schmidt, J. M., R. T. Good, B. Appleton, J. Sherrard, G. C. Raymant, M. R. Bogwitz, J. Martin, P. J. Daborn, M. E. Goddard, P. Batterham, et al. 2010. Copy number variation and transposable elements feature in recent, ongoing adaptation at the cyp6g1 locus. PLoS genetics, 6:e1000998.

Sellis, D., B. J. Callahan, D. A. Petrov, and P. W. Messer. 2011. Heterozygote advantage as a natural consequence of adaptation in diploids. Proceedings of the National Academy of Sciences, 108:20666–20671.

Siljestam, M., G. Arnqvist, and C. Rueffler. 2026. Sex-specific dominance maintains multilocus polymorphism and can resolve sexual conflict. bioRxiv.

Siljestam, M., and C. Rueffler. 2024. Heterozygote advantage can explain the extraordinary diversity of immune genes. eLife, 13:e94587.

Sinclair-Waters, M., T. Nome, J. Wang, S. Lien, M. P. Kent, H. Sægrov, B. Florø-Larsen, G. H. Bolstad, C. R. Primmer, and N. J. Barson. 2022. Dissecting the loci underlying maturation timing in atlantic salmon using haplotype and multi-snp based association methods. Heredity, 129:356–365.

Singh, A., and A. F. Agrawal. 2023. Two forms of sexual dimorphism in gene expression in drosophila melanogaster: their coincidence and evolutionary genetics. Molecular Biology and Evolution, 40:msad091.

Spencer, H. G., and C. Mitchell. 2016. The selective maintenance of allelic variation under generalized dominance. G3: Genes, Genomes, Genetics, 6:3725–3732.

Spencer, H. G., and N. K. Priest. 2016. The evolution of sex-specific dominance in response to sexually antagonistic selection. The American Naturalist, 187:658–666.

Telonis-Scott, M., A. Kopp, M. L. Wayne, S. V. Nuzhdin, and L. M. McIntyre. 2009. Sex-specific splicing in drosophila: widespread occurrence, tissue specificity and evolutionary conservation. Genetics, 181:421–434.

Tosto, N. M., E. R. Beasley, B. B. Wong, J. E. Mank, and S. P. Flanagan. 2023. The roles of sexual selection and sexual conflict in shaping patterns of genome and transcriptome variation. Nature Ecology & Evolution, 7:981–993.

Van Dooren, T. J. 1999. The evolutionary ecology of dominancerecessivity. Journal of theoretical Biology, 198:519–532.

Van Doorn, G. S., and U. Dieckmann. 2006. The long-term evolution of multilocus traits under frequency-dependent disruptive selection. Evolution, 60:2226–2238.

Wagstaff, B. J., and D. J. Begun. 2005. Molecular population genetics of accessory gland protein genes and testis-expressed genes in drosophila mojavensis and d. arizonae. Genetics, 171:1083–1101.

Wayne, M. L., M. Telonis-Scott, L. M. Bono, L. Harshman, A. Kopp,S. V. Nuzhdin, and L. M. McIntyre. 2007. Simpler mode of inheritance of transcriptional variation in male drosophila melanogaster. Proceedings of the National Academy of Sciences, 104:18577–18582.

Wilder, S. M., D. Raubenheimer, and S. J. Simpson. 2016. Moving beyond body condition indices as an estimate of fitness in ecological and evolutionary studies. Functional Ecology, 30:108–115.

Wittmann, M. J., A. O. Bergland, M. W. Feldman, P. S. Schmidt, andD. A. Petrov. 2017. Seasonally fluctuating selection can maintain polymorphism at many loci via segregation lift. Proceedings of the National Academy of Sciences, 114:E9932–E9941.

Wong, H. W., and L. Holman. 2023. Pleiotropic fitness effects across sexes and ages in the drosophila genome and transcriptome. Evolution, 77:2642–2655.

Wright, A. E., M. Fumagalli, C. R. Cooney, N. I. Bloch, F. G. Vieira,S. D. Buechel, N. Kolm, and J. E. Mank. 2018. Male-biased gene expression resolves sexual conflict through the evolution of sex-specific genetic architecture. Evolution Letters, 2:52–61.

Wright, S. 1931. Evolution in Mendelian populations. Genetics, 16:97–159.

Yeaman, S. 2022. Evolution of polygenic traits under global vs local adaptation. Genetics, 220:iyab134.

Zhou, Y., F. He, W. Pu, X. Gu, J. Wang, and Z. Su. 2020. The impact of dna methylation dynamics on the mutation rate during human germline development. G3: Genes, Genomes, Genetics, 10:3337–3346.

## References

Champagnat, N., R. Ferrière, and S. Méléard. 2006. Unifying evolutionary dynamics: from individual stochastic processes to macroscopic models. Theoretical Population Biology, 69:297–321.

Charnov, E. L. 1982. The Theory of Sex Allocation, vol. 18. Princeton University Press.

Dercole, F., and S. Rinaldi. 2008. Analysis of evolutionary processes: the adaptive dynamics approach and its applications: the adaptive dynamics approach and its applications. Princeton University Press.

Dieckmann, U., and M. Doebeli. 1999. On the origin of species by sympatric speciation. Nature, 400:354–357.

Doebeli, M. 2011. Adaptive Diversification (MPB-48). Princeton University Press.

Fisher, R. A. 1930. The genetical theory of natural selection. Oxford University Press.

Geritz, S. A. H., É. Kisdi, G. Meszéna, and J. A. J. Metz. 1998. Evolutionarily singular strategies and the adaptive growth and branching of the evolutionary tree. Evolutionary Ecology, 12:35– 57.

Leimar, O. 2001. Evolutionary change and darwinian demons. Selection, 2:65–72.

Levins, R. 1968. Evolution in Changing Environments: Some Theoretical Explorations. (MPB-2). Princeton University Press, Princeton, New Jersey.

Metz, J. A. J. 2008. Fitness. Pages 1599–1612 in S. E. Jorgensen and B. Fath, eds. Encyclopedia of ecology. Oxford: Elsevier, available online as IIASA Interim Report IR-06-061.

Priklopil, T., and L. Lehmann. 2020. Invasion implies substitution in ecological communities with class-structured populations. Theoretical population biology, 134:36–52.

Rueffler, C., T. J. Van Dooren, and J. A. Metz. 2004. Adaptive walks on changing landscapes: Levins’ approach extended. Theoretical population biology, 65:165–178.

Schmid, M., C. Rueffler, L. Lehmann, and C. Mullon. 2024. Resource variation within and between patches: Where exploitation competition, local adaptation, and kin selection meet. The American Naturalist, 203:E19–E34.

Shaw, R. F., and J. D. Mohler. 1953. The selective significance of the sex ratio. The American Naturalist, 87:337–342.

Slatkin, M. 1979. Frequency- and density-dependent selection on a quantitative trait. Genetics, 93:755–771.

Svardal, H., C. Rueffler, and J. Hermisson. 2015. A general condition for adaptive genetic polymorphism in temporally and spatially heterogeneous environments. Theoretical Population Biology, 99:76– 97.

Wright, S. 1931. Evolution in Mendelian populations. Genetics, 16:97–159.

